# The Tabula Sapiens: a multiple organ single cell transcriptomic atlas of humans

**DOI:** 10.1101/2021.07.19.452956

**Authors:** The Tabula Sapiens Consortium, Stephen R Quake

## Abstract

Molecular characterization of cell types using single cell transcriptome sequencing is revolutionizing cell biology and enabling new insights into the physiology of human organs. We created a human reference atlas comprising nearly 500,000 cells from 24 different tissues and organs, many from the same donor. This atlas enabled molecular characterization of more than 400 cell types, their distribution across tissues and tissue specific variation in gene expression. Using multiple tissues from a single donor enabled identification of the clonal distribution of T cells between tissues, the tissue specific mutation rate in B cells, and analysis of the cell cycle state and proliferative potential of shared cell types across tissues. Cell type specific RNA splicing was discovered and analyzed across tissues within an individual.

## Introduction

Although the genome is often called the blueprint of an organism, it is perhaps more accurate to describe it as a parts list composed of the various genes which may or may not be used in the different cell types of a multicellular organism. While nearly every cell in the body has essentially the same genome, each cell type makes different use of that genome and expresses a subset of all possible genes (*1*). Therefore, the genome in and of itself does not provide an understanding of the molecular complexity of the various cell types of that organism. This has motivated efforts to characterize the molecular composition of various cell types within humans and multiple model organisms, both by transcriptional (*2*) and proteomic (*3, 4*) approaches.

While such efforts are yielding insights (*5–7*), one caveat to current approaches is that individual organs are often collected at different locations, from different donors (*8*) and processed using different protocols, or lack replicate data (*9*). Controlled comparisons of cell types between different tissues and organs are especially difficult when donors differ in genetic background, age, environmental exposure, and epigenetic effects. To address this, we developed an approach to analyzing large numbers of organs from the same individual (*10*), which we originally used to characterize age-related changes in gene expression in various cell types in the mouse (*11*).

### Data Collection and Cell Type Representation

We collected multiple tissues from individual human donors (designated TSP 1-15) and performed coordinated single cell transcriptome analysis on live cells (*12*). We collected 17 tissues from one donor, 14 tissues from a second donor, and 5 tissues from two other donors (**Fig. 1**). We also collected smaller numbers of tissues from a further 11 donors, creating biological replicates for nearly all tissues. The donors comprise a range of ethnicities, are balanced by gender, have a mean age of 51 years and a variety of medical backgrounds (**table S1**). Single cell transcriptome sequencing was performed with both FACS sorted cells in well plates with smartseq2 amplification as well as 10x microfluidic droplet capture and amplification for each tissue (**fig. S1**). Tissue experts used a defined cell ontology terminology to annotate cell types consistently across the different tissues (*13*), leading to a total of 475 distinct cell types with reference transcriptome profiles (**tables S2, S3**). The full dataset can be explored online with the cellxgene tool via the Tabula Sapiens data portal (*14*).

**Figure 1.**
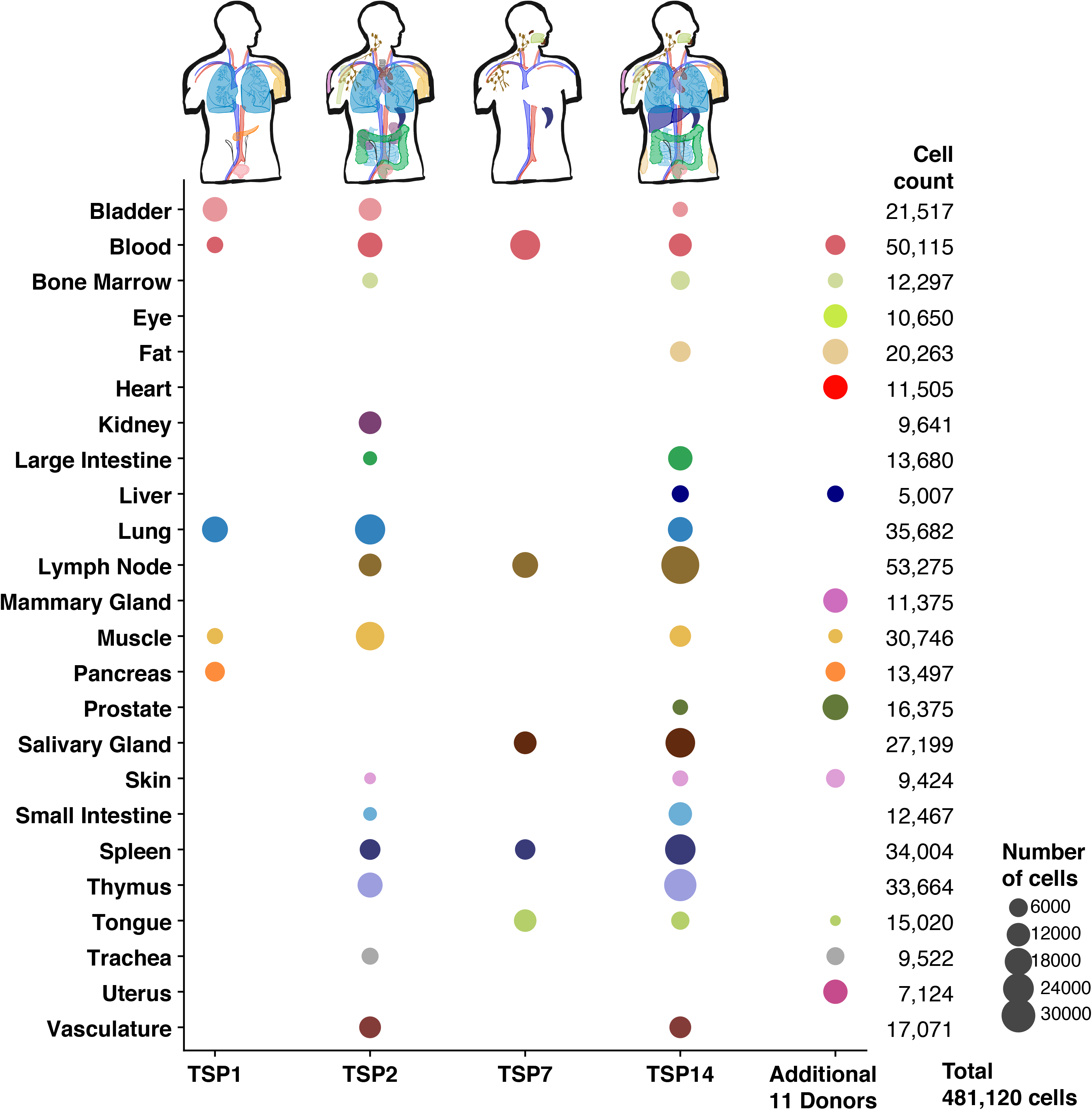
Overview of Tabula Sapiens. The Tabula Sapiens was constructed with data from 15 human donors; for detailed information on which tissues were examined for each donor please refer to **table S2**. Demographic and clinical information about each donor is listed in the supplement and in **table S1**. Donors 1, 2, 7 and 14 contributed the largest number of tissues each, and the number of cells from each tissue is indicated by the size of each circle. Tissue contributions from additional donors who contributed single or small numbers of tissues are shown in the “Additional donors” column, and the total number of cells for each organ are shown in the final column.

Data was collected for bladder, blood, bone marrow, eye, fat, heart, kidney, large intestine, liver, lung, lymph node, mammary, muscle, pancreas, prostate, salivary gland, skin, small intestine, spleen, thymus, tongue, trachea, uterus and vasculature. Fifty-nine separate specimens in total were collected, processed, and analyzed, and 481,120 cells passed QC filtering (**figs. S2-S7** and **table S2**). On a per compartment basis, the dataset includes 264,009 immune cells, 102,580 epithelial cells, 32,701 endothelial cells and 81,529 stromal cells. Working with live cells as opposed to isolated nuclei ensured that the dataset includes all mRNA transcripts within the cell, including transcripts that have been processed by the cell’s splicing machinery, thereby enabling insight into variation in alternative splicing.

To characterize the relationship between transcriptome data and conventional histologic analysis, a team of pathologists analyzed H&E stained sections prepared from 9 tissues from donor TSP2 and 13 tissues from donor TSP14 (*14*). Cells were identified by morphology and classified broadly into epithelial, endothelial, immune and stromal compartments, as well as rarely detected peripheral nervous system (PNS) cell types. (**Fig. 2A**). These classifications were used to estimate the relative abundances of cell types across four compartments, as well as to the uncertainties in these abundances due to spatial heterogeneity of each tissue type (**Fig. 2B, fig. S8**). We compared the histologically determined abundances with those obtained by single cell sequencing (**fig. S9**). Although, as expected, there can be substantial variation between the abundances determined by these methods, in aggregate we observe broad concordance over a large range of tissues and relative abundances. This approach enables an estimate of true cell type proportions since not every cell type survives dissociation with equal efficiency (*15*). For several of the tissues we also performed literature searches and collected tables of prior knowledge of cell type identity and abundance within those tissues (**table S4**). We compared literature values with our experimentally observed frequencies for three well annotated tissues: lung, muscle and bladder (**fig. S10**).

**Figure 2.**
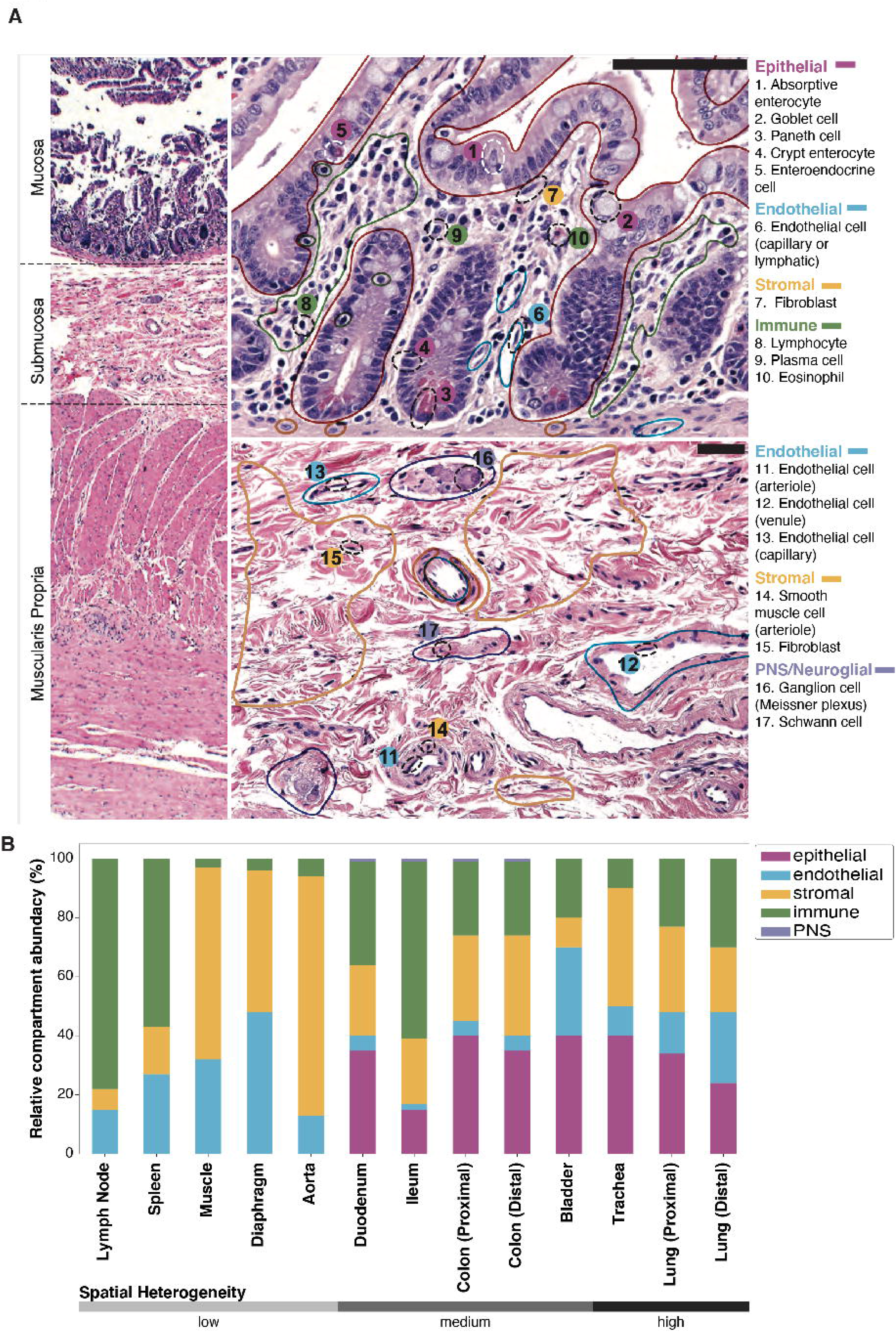
Comparison of single cell transcriptomics with conventional histology. Clinical pathology was performed on nine tissues from donors TSP2 and TSP13. **A**. Hematoxylin and eosin (H&E) stained image used for histology of the colon from TSP2, with compartments (solid, colored lines) and individual cell types (dashed black ellipses) identified by the pathologists. **B**. Coarse cell type representation of TSP2 as morphologically estimated by pathologists across several tissues, ordered by increasing heterogeneity of the tissue. Compartment colors are consistent between panels A and B.

### Immune Cells: Variation in Gene Expression Across Tissues and a Shared Lineage History

The Tabula Sapiens can be used to study differences in the gene expression programs and lineage histories of cell types that are shared across tissues. We analyzed tissue differences in the 36,475 macrophages distributed amongst 20 tissues, as tissue-resident macrophages are known to carry out specialized functions (*16*). These shared and orthogonal signatures are summarized in a correlation map (**fig. S11A**). For example, macrophages in the spleen were different from most other macrophages, and this was driven largely by higher expression of CD5L, a regulator of lipid synthesis (**fig. S11B**). We also observed a shared signature of elevated EREG (epiregulin) expression in solid tissues such as the skin, uterus and mammary compared to circulatory tissues (**fig. S11B**).

We characterized lineage relationships between T cells by assembling the T cell receptor sequences from from donor TSP2. Multiple T cell lineages were distributed across various tissues in the body, and we mapped their relationships (**Fig. 3A**). Large clones often reside in multiple organs, and several clones of mucosal associated invariant T cells are shared across donors (**fig. S11C**); these cells had characteristic expression of *TRAV1-2* as they are thought to be innate-like effector cells (*17*). Lineage information can also reveal tissue-specific somatic hyper-mutation rates in B cells. We assembled the B Cell Receptor sequences from donor TSP2 and inferred the germline ancestor of each cell. The mutational load varies dramatically by tissue of residence, with blood having the lowest mutational load compared to solid tissues (**fig. S11D**); solid tissues have an order of magnitude more mutations per nucleotide (mean=0.076, s.d.=0.026) compared to the blood (0.0069), suggesting that the immune infiltrates of solid tissues are dominated by mature B cells.

**Figure 3.**
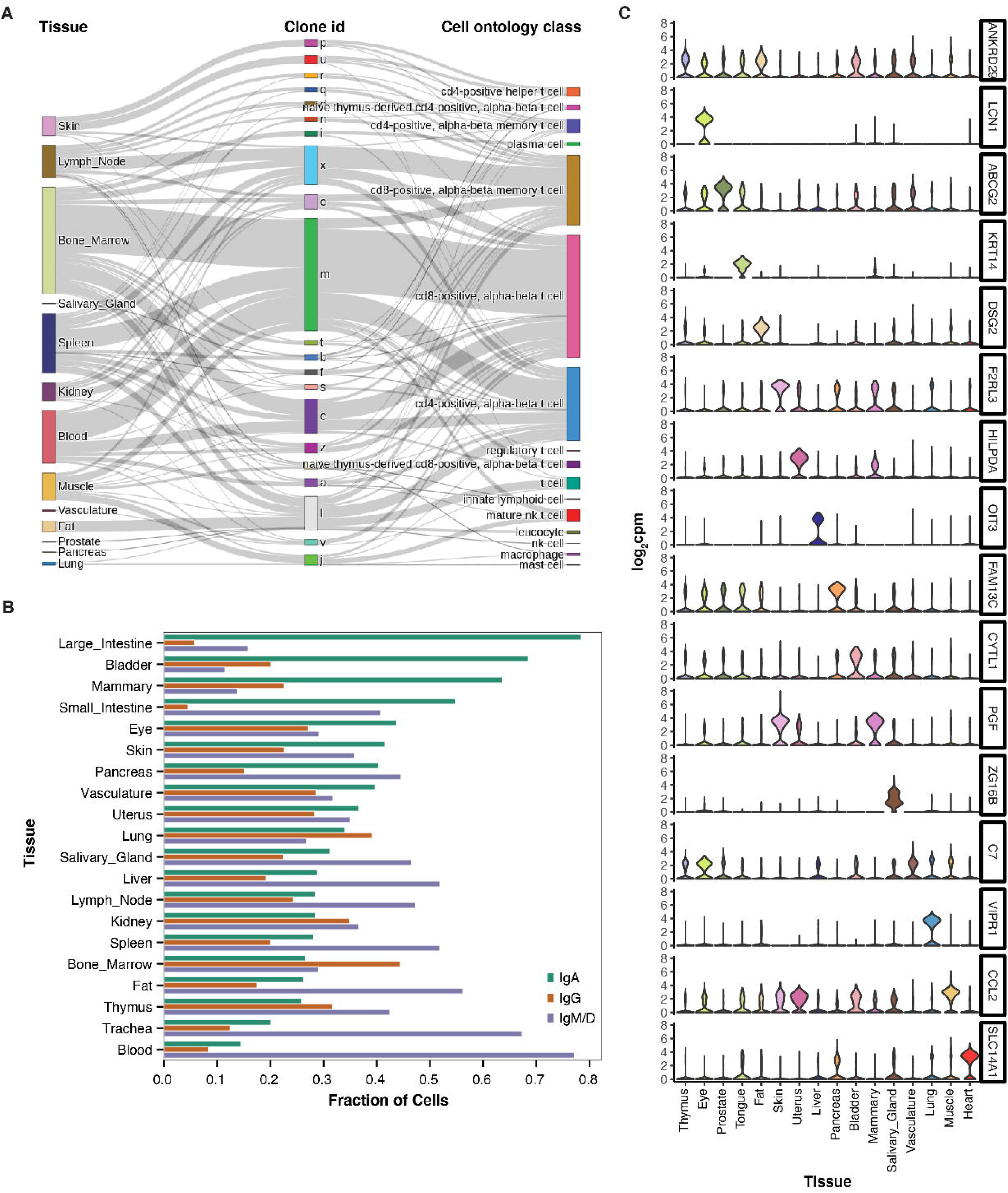
Analysis of immune and endothelial cell types shared across tissues. **A.** Illustration of clonal distribution of T cells across multiple tissues. The majority of T cell clones are found in multiple tissues and represent a variety of T cell subtypes. **B.** Prevalence of B cell isotypes across tissues, ordered by decreasing abundance of IgA. **C.** Expression level of tissue specific endothelial markers, shown as violin plots, in the entire dataset. Many of the markers are highly tissue specific, and typically derived from multiple donors as follows: bladder (3 donors), eye (2), fat (2), heart (1), liver (2), lung (3), mammary (1), muscle (4), pancreas (2), prostate (2), salivary gland (2), skin (2), thymus (2), tongue (2), uterus (1) and vasculature (2). A detailed donor-tissue breakdown is available in **table S2**.

B cells also undergo class-switch recombination which diversifies the humoral immune response by using constant region genes with distinct roles in immunity. We classified every B cell in the dataset as IgA, IgG, or IgM expressing and then calculated the relative amounts of each cellular isotype in each tissue (**Fig. 3B, table S5**). Secretory IgA is known to interact with pathogens and commensals at the mucosae, IgG is often involved in direct neutralization of pathogens, and IgM is typically expressed in naive B cells or secreted in first response to pathogens. Consistent with this, our analysis revealed opposing gradients of prevalence of IgA and IgM expressing B cells across the tissues with blood having the lowest relative abundance of IgA producing cells and the large intestine having the highest relative abundance, and the converse for IgM expressing B cells (**Fig. 3B**).

### Endothelial Cells Subtypes with Tissue-Specific Gene Expression Programs

As another example of analyzing shared cell types across organs, we focused on endothelial cells (ECs). While ECs are often categorized as a single cell type, they exhibit differences in morphology, structure, immunomodulatory and metabolic phenotypes depending on their tissue of origin. Here, we discovered that tissue-specificity is also reflected in their transcriptomes, as ECs mainly cluster by tissue-of-origin (**table S6**). UMAP analysis (**fig. S12A**) revealed that lung, heart, uterus, liver, pancreas, fat and muscle ECs exhibited the most distinct transcriptional signatures, reflecting their highly specialized roles. These distributions were conserved across donors (**fig. S12B**).

Interestingly, ECs from the thymus, vasculature, prostate, and eye were similarly distributed across several clusters, suggesting not only similarity in transcriptional profiles but in their sources of heterogeneity. Differential gene expression analysis between ECs of these 16 tissues revealed several canonical and previously undescribed tissue-specific vascular markers (**Fig. 3C**). These data recapitulate tissue-specific vascular markers such as LCN1 (tear lipocalin) in the eye, ABCG2 (transporter at the blood-testis barrier) in the prostate, and OIT3 (oncoprotein induced transcript 3) in the liver. Of the potential novel markers determined by this analysis, SLC14A1 (solute carrier family 14 member 1) appears to be a new specific marker for endothelial cells in the heart, whose expression was independently validated with data from the Human Protein Atlas (*18*) (**fig. S13**).

Notably, lung ECs formed two distinct populations, which is in line with the aerocyte (aCap-EDNRB+) and general capillary (gCap - PLVAP+) cells described in the mouse and human lung (*19*) (**fig. S12 C,D**). The transcriptional profile of gCaps were also more similar to ECs from other tissues, indicative of their general vascular functions in contrast to the more specialized aCap populations. Lastly, we detected two distinct populations of ECs in the muscle, including a MSX1+ population with strong angiogenic and endothelial cell proliferation signatures, and a CYP1B1+ population enriched in metabolic genes, suggesting the presence of functional specialization in the muscle vasculature (**fig. S12 E,F**).

### Alternative Splice Variants are Cell Type Specific

We used SICILIAN (*20*) to identify alternative splice junctions in Tabula Sapiens using both 10X and Smart-Seq2 sequencing technologies and found a total of 955,785 junctions (**fig. S14A-E, table S7**). 217,855 of these were previously annotated, and thus our data provides independent validation of 61% of the total junctions catalogued in the entire RefSeq database. Although annotated junctions made up only 22.8% of the unique junctions, they represent 93% of total reads, indicating that previously annotated junctions tend to be expressed at higher levels than novel junctions. We additionally found 34,624 novel junctions between previously annotated 3’ and 5’ splice sites (3.6%). We also identified 119,276 junctions between a previously annotated site and a novel site in the gene (12.4%). This leaves 584,030 putative junctions for which both splice sites were previously unannotated, i.e. about 61% of the total detected junctions. Most of these have at least one end in a known gene (94.7%), while the remainder represent potential new splice variants from unannotated regions (5.3%). In the absence of independent validation, we conservatively characterized all of the unannotated splices as putative novel junctions. We then used the GTEx database (*21*) to seek independent corroborating evidence of these putative novel junctions, and found that reads corresponding to nearly one third of these novel junctions can be found within GTEx data (**table S7**); this corresponds to more than 300,000 new validated splice variants revealed by the Tabula Sapiens.

Hundreds of splice variants are used in a highly cell-type specific fashion; these can be explored in the cellxgene browser (*14*) which uses a statistic called SpliZ (*22*). Here we focus on two examples of cell type specific splicing of two well studied genes: MYL6 and CD47; similar cell-type specific splice usage was also observed with TPM1, TPM2, and ATP5F1C, three other genes with well-characterized splice variants (**fig. S15**).

MYL6 is an “essential light chain” (ELC) for myosin and is highly expressed in all tissues and compartments. Yet, splicing of MYL6, in particular involving the inclusion/exclusion of exon 6 (**Fig. 4A**) varies in a cell-type and compartment-specific manner (**Fig. 4B**). While the -exon6 isoform has previously been mainly described in phasic smooth muscle (*23*), we discovered it can also be the predominant isoform in non-smooth-muscle cell types. Our analysis establishes pervasive regulation of MYL6 splicing in many cell types, such as endothelial and immune cells. These previously unknown compartment-specific expression patterns of the two MYL6 isoforms are reproduced in multiple individuals from the Tabula Sapiens dataset (**Fig. 4A,B**).

**Figure 4.**
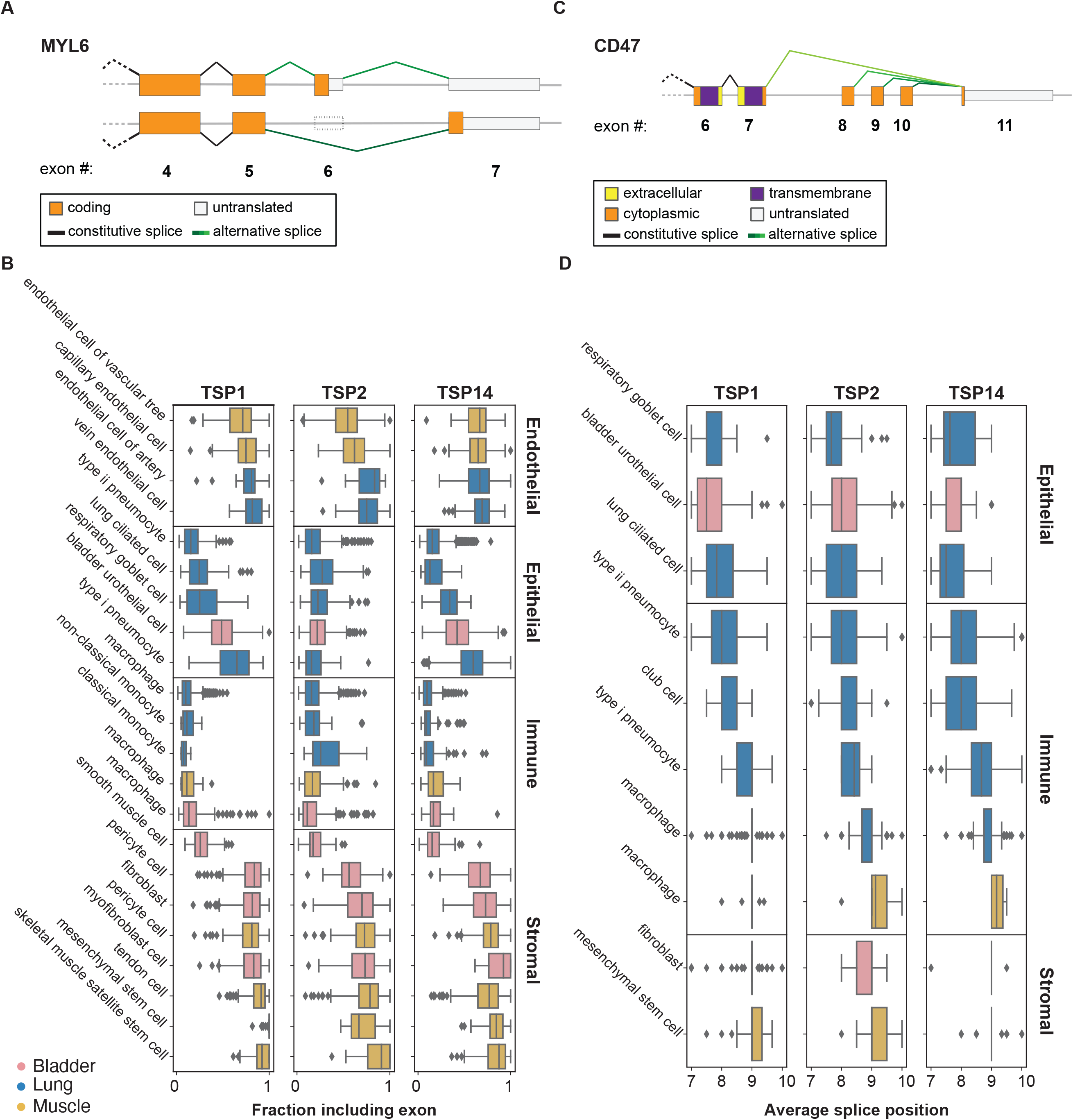
Alternative splicing analysis. **A,B**. The sixth exon in MYL6 is skipped at different proportions in different compartments. Cells in the immune and epithelial compartments tend to skip the exon, whereas cells in the endothelial and stromal compartments tend to include the exon. Boxes are grouped by compartment and colored by tissue. The fraction of junctional reads that include exon 6 was calculated for each cell with more than 10 reads mapping to the exon skipping event. Horizontal box plots in **B** show the distribution of exon inclusion in each cell type. **C,D**. Alternative splicing in CD47 involves one 5’ splice site (exon 11, 108,047,292) and four 3’ splice sites. Horizontal box plots in **D** show the distribution of weighted averages of alternative 3’ splice sites in each cell type. Epithelial cells tend to use closer exons to the 5’ splice site compared to immune and stromal cells. Boxes are grouped by compartment and colored by tissue.

CD47 is a multi-spanning membrane protein involved in many cellular processes, including angiogenesis, cell migration, and as a “don’t eat me” signal to macrophages (*24*). Differential use of exons 7-10 (**Fig. 4C and fig. S14F**) compose a variably long cytoplasmic tail (*25*). Immune cells – but also stromal and endothelial cells – have a distinct, consistent splicing pattern in CD47 that dominantly excludes two proximal exons and splicing directly to exon 8. In contrast to other compartments, epithelial cells exhibit a different splicing pattern that increases the length of the cytoplasmic tail by splicing more commonly to exon 9 and exon 10 (**Fig. 4D**). Characterization of the splicing programs of CD47 in single cells may have implications for understanding the differential signaling activities of CD47 and for therapeutic manipulation of CD47 function.

### Cell State Dynamics Can Be Inferred From A Single Time Point

Although the Tabula Sapiens was created from a single moment in time for each donor, it is possible to infer dynamic information from the data. Cell division is an important transient change of internal cell state, and we computed a cycling index for each cell type to identify actively proliferating versus quiescent or post-mitotic cell states. Rapidly dividing progenitor cells had among the highest cycling indices, while cell types from the endothelial and stromal compartments, which are known to be largely quiescent, had low cycling indices (**Fig. 5A**). In intestinal tissue, transient amplifying cells and the crypt stem cells divide rapidly in the intestinal crypts to give rise to terminally differentiated cell types of the villi (*26*). These cells were ranked with the highest cycling indices whereas terminally differentiated cell types such as the goblet cells had the lowest ranks (**fig. S16A**). To complement the computational analysis of cell cycling, we performed immunostaining of intestinal tissue for MKI67 protein (commonly referred to as Ki-67) and confirmed that transient amplifying cells abundantly express this proliferation marker (**fig. S16B,C**), supporting that this marker is differentially expressed in the G2/M cluster.

**Figure 5.**
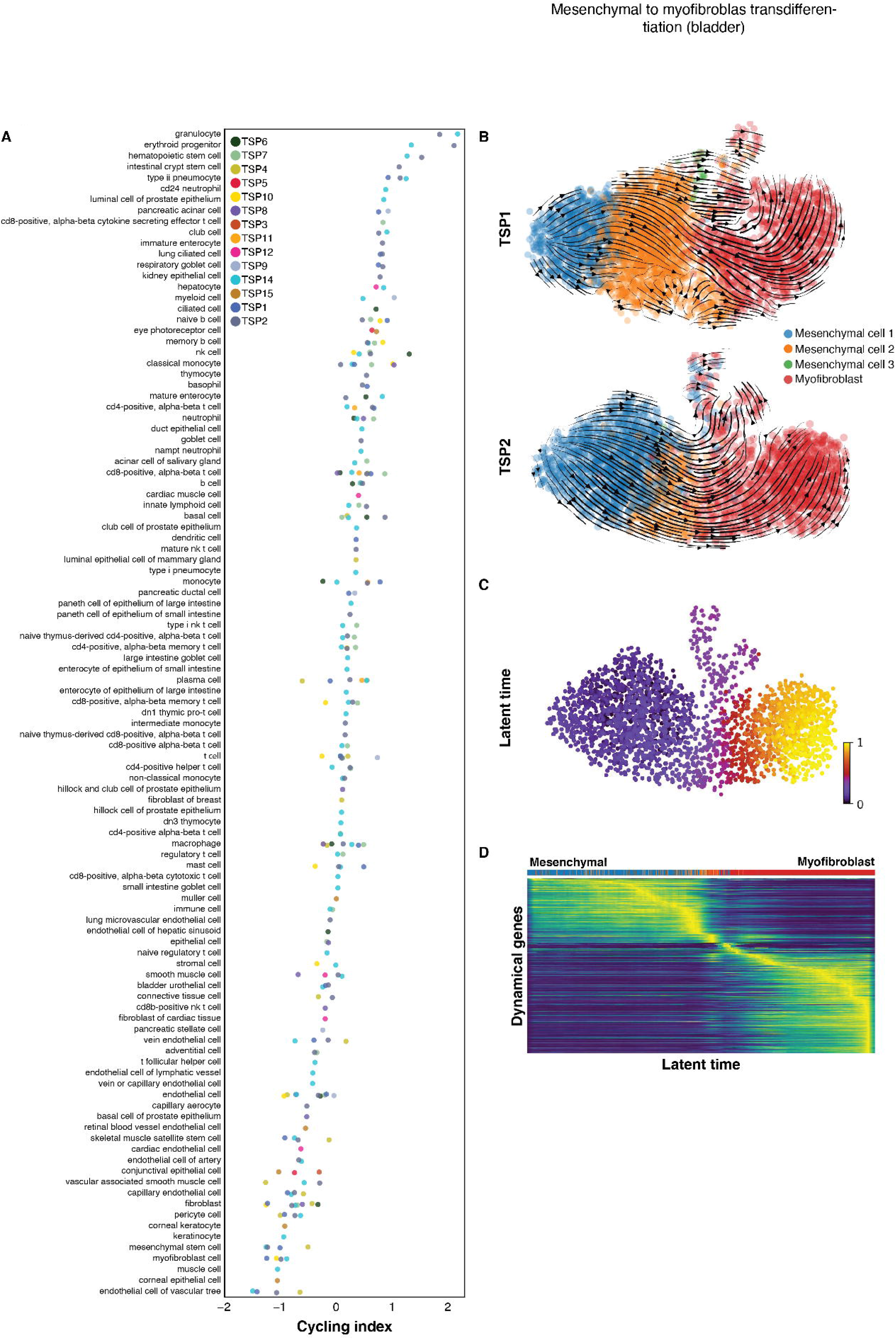
Dynamic changes in cell state. **A**. Cell types ordered by magnitude of cell cycling index, per donor (each a separate color) with the most highly proliferative at the top and quiescent cells at the bottom of the list. **B**. RNA velocity analysis demonstrating mesenchymal to myofibroblast transition in the bladder. The arrows represent a flow derived from the ratio of unspliced to spliced transcripts which in turn predicts dynamic changes in cell identity. **C**,**D.** Latent time analysis of the mesenchymal to myofibroblast transition in the bladder demonstrating stereotyped changes in gene expression trajectory.

We observed several interesting tissue-specific differences in cell cycling. To illustrate one example, UMAP clustering of macrophages showed tissue-specific clustering of this cell type, and that blood, bone marrow, and lung macrophages have the highest cycling indices compared to macrophages found in the bladder, skin, and muscle (**fig. S16D-G**). Consistent with this finding, the expression values of CDK-inhibitors (in particular the gene CDKN1A), which block the cell cycle, have the lowest overall expression in macrophages from tissues with high cycling indices (**fig. S16F**).

We used RNA velocity (*27*) as a further dynamic approach to study trans-differentiation of bladder mesenchymal cells to myofibroblasts (**Fig. 5B**). Latent time analysis, which provides an estimate of each cell’s internal clock using RNA velocity trajectories (*28*), correctly identified the direction of differentiation (**Fig. 5C**) across multiple donors. Ordering cells as a function of latent time shows clustering of the mesenchymal and myofibroblast gene expression programs for the most dynamically expressed genes (**Fig. 5D**). Among these genes, ACTN1 (Alpha Actinin 1) – a key actin crosslinking protein that stabilizes cytoskeleton-membrane interactions (*29*) – increases across the mesenchymal to myofibroblast trans-differentiation trajectory (**fig. S16H**). Another gene with a similar trajectory is MYLK (myosin light-chain kinase), which also rises as myofibroblasts attain more muscle-like properties (*30*). Finally, a random sampling of the most dynamic genes shared across TSP1 and TSP2 demonstrated that they share concordant trajectories and revealed some of the core genes in the transcriptional program underlying this trans-differentiation event within the bladder (**fig. S16I**).

### Unexpected Spatial Variation in the Microbiome

The Tabula Sapiens provided an opportunity to densely and directly sample the human microbiome throughout the gastrointestinal tract. The intestines from donors TSP2 and TSP14 were sectioned into five regions: the duodenum, jejunum, ileum, and ascending and sigmoid colon (**Fig. 6A**). Each section was transected, and three to nine samples of were collected from each location, followed by amplification and sequencing of the 16S rRNA gene. Uniformly there was a high (∼10-30%) relative abundance of Proteobacteria, particularly Enterobacteriaceae (**Fig. 6B**), even in the colon. Samples from each of the duodenum, jejunum, and ileum were largely distinct from one another, with samples exhibiting individual patterns of blooming or absence of certain families (**Fig. 6B**). These data reveal that the microbiota is patchy even at a 3-inch length scale. We observed similar heterogeneity in both donors (**fig. S17A-C**). In the small intestine, richness (number of observed species) was also variable, and was negatively correlated with the relative abundance of Burkholderiaceae (**Fig. 6B**); in TSP2, the Proteobacteria phylum was dominated by Enterobacteriaceae, which was present at >30% in all samples at a level negatively correlated with richness (**fig. S17A-C**). In a comparison of species from adjacent regions across the gut, a large fraction of species was unique to each region (**Fig. 6C**), reflecting the patchiness. These data are derived from only two donor samples and further conclusions about the statistics and extent of microbial patchiness will require larger studies.

**Figure 6.**
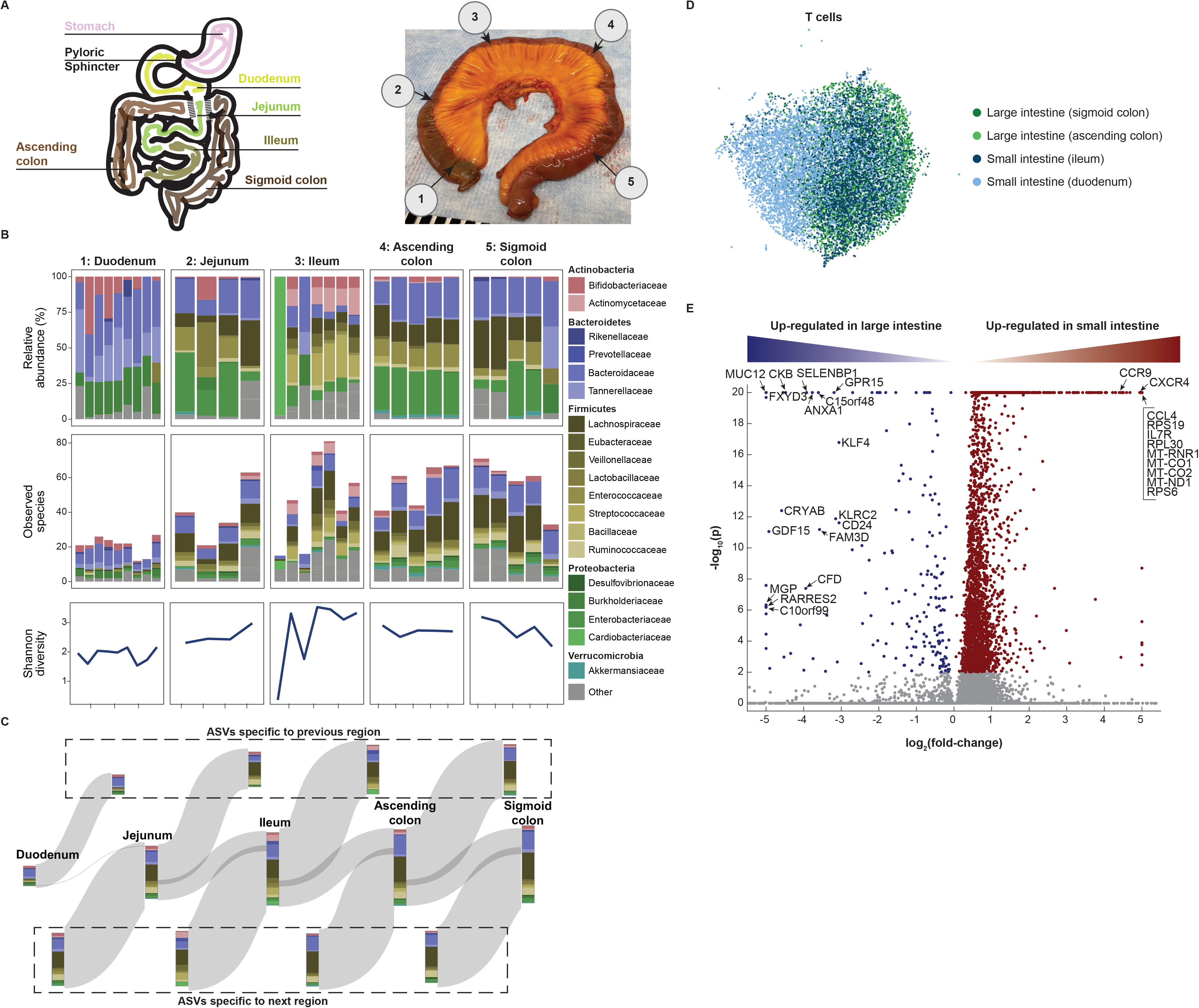
High-resolution view highlights patchiness of the gut microbiome. **A**. Schematic (left) and photo of the colon from donor TSP2 (right), with numbers 1-5 representing microbiota sampling locations. **B**. Relative abundances and richness (number of observed species) at the family level in each sampling location, as determined by 16S rRNA sequencing. The Shannon diversity, a metric of evenness, mimics richness. Variability in relative abundance and/or richness/Shannon diversity was higher in the duodenum, jejunum and ileum as compared with the ascending and sigmoid colon. **C**. A Sankey diagram showing the inflow and outflow of microbial species from each section of the gastrointestinal tract. The stacked bar for each gastrointestinal section represents the number of observed species in each family as the union of all sampling locations for that section. The stacked bar flowing out represents gastrointestinal species not found in the subsequent section and the stacked bar flowing into each gastrointestinal section represents the species not found in the previous section. **D**. UMAP clustering of T cells reveals distinct transcriptome profiles in the distal and proximal small and large intestines. **E**. Dots in volcano plot highlight genes up-regulated in the large (left) and small (right) intestines. Labeled dots include genes with known roles in trafficking, survival, and activation.

We analyzed host immune cells in conjunction with the spatial microbiome data; UMAP clustering analysis revealed that the small intestine T cell pool from TSP14 contained a population with distinct transcriptomes (**Fig. 6D**). The most significant transcriptional differences in T cells between the small and large intestine were genes associated with trafficking, survival, and activation (**Fig. 6E**, **table S8**). For example, expression of the long non-coding RNA MALAT1, which impacts the regulatory function of T cells, and CCR9, which mediates T lymphocyte development and migration to the intestine (*31*), were high only in the small intestine, while GPR15 (colonic T cell trafficking), SELENBP1 (selenium transporter), ANXA1 (repressor of inflammation in T cells), KLRC2 (T cell lectin), CD24 (T cell survival), GDF15 (T cell inhibitor), and RARRES2 (T cell chemokine) exhibited much higher expression in the large intestine. Within the epithelial cells, we observed distinct transcriptomes between small and large intestine Paneth cells and between small and large intestine enterocytes, while there was some degree of overlap for each of the two cell types for either location (**fig. S17E,F**). The site-specific composition of the microbiome in the intestine, paired with distinct T cell populations at each site helps define local host-microbe interactions that occur in the GI tract and is likely reflective of a gradient of physiological conditions that influence host-microbe dynamics.

## Conclusion

The Tabula Sapiens is part of a growing set of data which when analyzed together will enable many interesting comparisons of both a biological and a technical nature. Studying particular cell types across organs, datasets, and species will yield new biological insights – as shown with fibroblasts (*32*). Similarly, comparing fetal human cell types (*33*) to those determined here in adults may give insight into the loss of plasticity from early development to maturity. Having multi-organ data from individual donors may facilitate development of methods to compare diverse datasets and yield understanding of technical artifacts from various approaches (*8, 9, 34, 35*). The Tabula Sapiens has enabled discoveries relating to shared behavior and organ specific differences across cell types. For example, we found T cell clones shared between organs, and characterized organ dependent hypermutation rates amongst resident B cells. Endothelial cells and macrophages are cell types which are shared across tissues, but often show subtle tissue-specific differences in gene expression. We found an unexpectedly large and diverse amount of cell-type specific RNA splice variant usage, and discovered and validated many new splices. These are but a few examples of how the Tabula Sapiens represents a broadly useful reference to understand and explore human biology deeply at cellular resolution.

### Brief synopsis of methods

Fresh whole non-transplantable organs, or 1-2cm^2^ organ samples, were obtained from surgery and then transported on ice by courier to tissue expert labs where they were immediately prepared for transcriptome sequencing. Single-cell suspensions were prepared for 10x Genomics 3’ V3.1 droplet-based sequencing and for FACS sorted 384-well plate Smart-seq2. Preparation began with dissection, digestion with enzymes and physical manipulation; tissue specific details are in the methods supplement (*12*). Cell suspensions from some organs were normalized by major cell compartment (epithelial, endothelial, immune, and stromal) using antibody-labelled magnetic microbeads to enrich rare cell types. cDNA and sequencing libraries were prepared and run on the Illumina NovaSeq 6000 with the goal to obtain 10,000 droplet-based cells and 1000 plate-based cells for each organ. Sequences were de-multiplexed and aligned to the GRCh38 reference genome. Gene count tables were generated with CellRanger (droplet samples), or STAR and HTSEQ (plate samples). Cells with low UMI counts and low gene counts were removed. Droplet cells were filtered to remove barcode-hopping events and filtered for ambient RNA using DecontX. Sequencing batches were harmonized using scVI and projected to 2-D space with UMAP for analysis by the tissue experts. Expert annotation was made through the cellxgene browser and regularized with a public cell ontology. Annotation was manually QC checked and cross-validated with PopV, an annotation tool, which employs seven different automated annotation methods. For complete methods, see supplementary materials (*12*).

## Acknowledgements

We express our gratitude and thanks to donor WEM and his family, as well as to all of the anonymous organ and tissue donors and their families for giving both the gift of life and the gift of knowledge by their generous donations. We also thank Donor Network West for their cooperation in this project, S. Schmid for a close reading of the manuscript, and B. Tojo for the original artwork in Figure 1. This project has been made possible in part by grant number 2019-203354 from the Chan Zuckerberg Initiative DAF, an advised fund of Silicon Valley Community Foundation, and by support from the Chan Zuckerberg Biohub. The data portal for this publication is available at (*14*).

## Data and materials availability

The entire dataset can be explored interactively at http://tabula-sapiens-portal.ds.czbiohub.org/(14). The code used for the analysis is available from Zenodo (https://doi.org/10.5281/zenodo.6069683) (*36*). Gene counts and metadata are available from figshare (https://doi.org/10.6084/m9.figshare.14267219) (*37*) and have been deposited in the Gene Expression Omnibus (GSE149590); the raw data files are available from a public AWS S3 bucket (https://registry.opendata.aws/tabula-sapiens/) and instructions on how to access the data have been provided in the project GitHub. The histology images are available from figshare (https://doi.org/10.6084/m9.figshare.14962947) (*38*). SpliZ scores are available from figshare (https://doi.org/10.6084/m9.figshare.14977281) (*39*).

## The Tabula Sapiens Consortium Author List

### Overall Project Direction and Coordination

Robert C. Jones^1^, Jim Karkanias^2^, Mark Krasnow^3,4^, Angela Oliveira Pisco^2^, Stephen R. Quake^1,2,5^, Julia Salzman^3,6^, Nir Yosef^2,7,8,9^

### Donor Recruitment

Bryan Bulthaup^10^, Phillip Brown^10^, William Harper^10^, Marisa Hemenez^10^, Ravikumar Ponnusamy^10^, Ahmad Salehi^10^, Bhavani A. Sanagavarapu^10^, Eileen Spallino^10^

### Surgeons

Ksenia A. Aaron^11^, Waldo Concepcion^10^, James M. Gardner^12,13^, Burnett Kelly^10,14^, Nikole Neidlinger^10^, Zifa Wang^10^

### Logistical coordination

Sheela Crasta^1,2^, Saroja Kolluru^1,2^, Maurizio Morri^2^, Angela Oliveira Pisco^2^, Serena Y. Tan^15^, Kyle J. Travaglini^3^, Chenling Xu^7^

### Organ Processing

Marcela Alcántara-Hernández^16^, Nicole Almanzar^17^, Jane Antony^18^, Benjamin Beyersdorf^19^, Deviana Burhan^20^, Kruti Calcuttawala^21^, Matthew M. Carter^16^, Charles K. F. Chan^18,22^, Charles A. Chang^23^, Stephen Chang^3,19^, Alex Colville^21,24^, Sheela Crasta^1,2^, Rebecca N. Culver^25^, Ivana Cvijović^1,5^, Gaetano D’Amato^26^, Camille Ezran^3^, Francisco X. Galdos^18^, Astrid Gillich^3^, William R. Goodyer^27^, Yan Hang^23,28^, Alyssa Hayashi^1^, Sahar Houshdaran^29^, Xianxi Huang^19,30^, Juan C. Irwin^29^, SoRi Jang^3^, Julia Vallve Juanico^29^, Aaron M. Kershner^18^, Soochi Kim^21,24^, Bernhard Kiss^18^, Saroja Kolluru^1,2^, William Kong^18^, Maya E. Kumar^17^, Angera H. Kuo^18^, Rebecca Leylek^16^, Baoxiang Li^31^, Gabriel B. Loeb^32^, Wan-Jin Lu^18^, Sruthi Mantri^33^, Maxim Markovic^1^, Patrick L. McAlpine^11,34^, Antoine de Morree^21,24^, Maurizio Morri^2^, Karim Mrouj^18^, Shravani Mukherjee^31^, Tyler Muser^17^, Patrick Neuhöfer^3,35,36^, Thi D. Nguyen^37^, Kimberly Perez^16^, Ragini Phansalkar^26^, Angela Oliveira Pisco^2^, Nazan Puluca^18^, Zhen Qi^18^, Poorvi Rao^20^, Hayley Raquer-McKay^16^, Nicholas Schaum^18,21^, Bronwyn Scott^31^, Bobak Seddighzadeh^38^, Joe Segal^20^, Sushmita Sen^29^, Shaheen Sikandar^18^, Sean P. Spencer^16^, Lea Steffes^17^, Varun R. Subramaniam^31^, Aditi Swarup^31^, Michael Swift^1,^ Kyle J. Travaglini^3^, Will Van Treuren^16^, Emily Trimm^26^, Stefan Veizades^19,39^, Sivakamasundari Vijayakumar^18^, Kim Chi Vo^29^, Sevahn K. Vorperian^1,40^, Wanxin Wang^29^, Hannah N.W. Weinstein^38^, Juliane Winkler^41^, Timothy T.H. Wu^3^, Jamie Xie^38^, Andrea R.Yung^3^, Yue Zhang^3^

### Sequencing

Angela M. Detweiler^2^, Honey Mekonen^2^, Norma F. Neff^2^, Rene V. Sit^2^, Michelle Tan^2^, Jia Yan^2^

### Histology

Gregory R. Bean^15^, Vivek Charu^15^, Erna Forgó^15^, Brock A. Martin^15^, Michael G. Ozawa^15^, Oscar Silva^15^, Serena Y. Tan^15^, Angus Toland^15^, Venkata N.P. Vemuri^2^

### Data Analysis

Shaked Afik^7^, Kyle Awayan^2^, Olga Borisovna Botvinnik^2^, Ashley Byrne^2^, Michelle Chen^1^, Roozbeh Dehghannasiri^3,6^, Angela M. Detweiler^2^, Adam Gayoso^7^, Alejandro A Granados^2^, Qiqing Li^2^, Gita Mahmoudabadi^1^, Aaron McGeever^2^, Antoine de Morree^21,24^, Julia Eve Olivieri^3,6,42^, Madeline Park^2^, Angela Oliveira Pisco^2^, Neha Ravikumar^1^, Julia Salzman^3,6^, Geoff Stanley^1^, Michael Swift^1^, Michelle Tan^2^, Weilun Tan^2^, Alexander J Tarashansky^2^, Rohan Vanheusden^2^, Sevahn K. Vorperian^1,40^, Peter Wang^3,6^, Sheng Wang^2^, Galen Xing^2^, Chenling Xu^6^, Nir Yosef^2,6,7,8^

### Expert Cell Type Annotation

Marcela Alcántara-Hernández^16^, Jane Antony^18^, Charles K. F. Chan^18,22^, Charles A. Chang^23^, Alex Colville^21,24^, Sheela Crasta^1,2^, Rebecca Culver^25^, Les Dethlefsen^43^, Camille Ezran^3^, Astrid Gillich^3^, Yan Hang^23,28^, Po-Yi Ho^16^, Juan C. Irwin^29^, SoRi Jang^3^, Aaron M. Kershner^18^, William Kong^18^, Maya E Kumar^17^, Angera H. Kuo^18^, Rebecca Leylek^16^, Shixuan Liu^3,44^, Gabriel B. Loeb^32^, Wan-Jin Lu^18^, Jonathan S Maltzman^45,46^, Ross J. Metzger^27,47^, Antoine de Morree^21,24^, Patrick Neuhöfer^3,35,36^, Kimberly Perez^16^, Ragini Phansalkar^26^, Zhen Qi^18^, Poorvi Rao^20^, Hayley Raquer-McKay^16,^ Koki Sasagawa^19,^ Bronwyn Scott^31^, Rahul Sinha^15,18,35^, Hanbing Song^38^, Sean P. Spencer^16^, Aditi Swarup^31^, Michael Swift^1^, Kyle J. Travaglini^3^, Emily Trimm^26^, Stefan Veizades^19,39^, Sivakamasundari Vijayakumar^18^, Bruce Wang^20^, Wanxin Wang^29^, Juliane Winkler^41^, Jamie Xie^38^, Andrea R.Yung^3^

### Tissue Expert Principal Investigators

Steven E. Artandi^3,35,36^, Philip A. Beachy^18,23,48^, Michael F. Clarke^18^, Linda C. Giudice^29^, Franklin W. Huang^38,49^, Kerwyn Casey Huang^1,16^, Juliana Idoyaga^16^, Seung K Kim^23,28^, Mark Krasnow^3,4^, Christin S. Kuo^17^, Patricia Nguyen^19,39,46^, Stephen R. Quake^1,2,5^, Thomas A. Rando^21,24^, Kristy Red-Horse^26^, Jeremy Reiter^50^, David A. Relman^16,43,46^, Justin L. Sonnenburg^16^, Bruce Wang^20^, Albert Wu^31^, Sean M. Wu^19,39^, Tony Wyss-Coray^21,24^

## Affiliations

^1^Department of Bioengineering, Stanford University; Stanford, CA, USA.

^2^Chan Zuckerberg Biohub; San Francisco, CA, USA.

^3^Department of Biochemistry, Stanford University School of Medicine; Stanford, CA, USA.

^4^Howard Hughes Medical Institute; USA.

^5^Department of Applied Physics, Stanford University; Stanford, CA, USA.

^6^Department of Biomedical Data Science, Stanford University; Stanford, CA, USA.

^7^Center for Computational Biology, University of California Berkeley; Berkeley, CA, USA.

^8^Department of Electrical Engineering and Computer Sciences, University of California Berkeley; Berkeley, CA, USA.

^9^Ragon Institute of MGH, MIT and Harvard; Cambridge, MA, USA.

^10^Donor Network West; San Ramon, CA, USA.

^11^Department of Otolaryngology-Head and Neck Surgery, Stanford University School of Medicine; Stanford, California, USA.

^12^Department of Surgery, University of California San Francisco; San Francisco, CA, USA.

^13^Diabetes Center, University of California San Francisco; San Francisco, CA, USA.

^14^DCI Donor Services; Sacramento, CA, USA.

^15^Department of Pathology, Stanford University School of Medicine; Stanford, CA, USA.

^16^Department of Microbiology and Immunology, Stanford University School of Medicine; Stanford, CA, USA.

^17^Department of Pediatrics, Division of Pulmonary Medicine, Stanford University; Stanford, CA, USA.

^18^Institute for Stem Cell Biology and Regenerative Medicine, Stanford University School of Medicine; Stanford, CA, USA.

^19^Department of Medicine, Division of Cardiovascular Medicine, Stanford University; Stanford, CA, USA.

^20^Department of Medicine and Liver Center, University of California San Francisco; San Francisco, CA, USA.

^21^Department of Neurology and Neurological Sciences, Stanford University School of Medicine; Stanford, CA, USA.

^22^Department of Surgery - Plastic and Reconstructive Surgery, Stanford University School of Medicine; Stanford, CA, USA.

^23^Department of Developmental Biology, Stanford University School of Medicine, Stanford, CA, USA

^24^Paul F. Glenn Center for the Biology of Aging, Stanford University School of Medicine; Stanford, CA, USA.

^25^Department of Genetics, Stanford University School of Medicine; Stanford, CA, USA.

^26^Department of Biology, Stanford University; Stanford, CA, USA.

^27^Department of Pediatrics, Division of Cardiology, Stanford University School of Medicine; Stanford, CA, USA.

^28^Stanford Diabetes Research Center, Stanford University School of Medicine, Stanford, California

^29^Center for Gynecology and Reproductive Sciences, Department of Obstetrics, Gynecology and Reproductive Sciences, University of California San Francisco; San Francisco, CA, USA.

^30^Department of Critical Care Medicine, The First Affiliated Hospital of Shantou University Medical College; Shantou, China.

^31^Department of Ophthalmology, Stanford University School of Medicine; Stanford, CA, USA.

^32^Division of Nephrology, Department of Medicine, University of California San Francisco; San Francisco, CA, USA.

^33^Stanford University School of Medicine; Stanford, CA, USA.

^34^Mass Spectrometry Platform, Chan Zuckerberg Biohub; Stanford, CA, USA.

^35^Stanford Cancer Institute, Stanford University School of Medicine; Stanford, CA, USA.

^36^Department of Medicine, Division of Hematology, Stanford University School of Medicine, Stanford, CA, USA

^37^Department of Biochemistry and Biophysics, Cardiovascular Research Institute, University of California San Francisco; San Francisco, CA, USA.

^38^Division of Hematology and Oncology, Department of Medicine, Bakar Computational Health Sciences Institute, Institute for Human Genetics, University of California San Francisco; San Francisco, CA, USA.

^39^Stanford Cardiovascular Institute; Stanford CA, USA.

^40^Department of Chemical Engineering, Stanford University; Stanford, CA, USA.

^41^Department of Cell & Tissue Biology, University of California San Francisco; San Francisco, CA, USA.

^42^Institute for Computational and Mathematical Engineering, Stanford University; Stanford, CA, USA.

^43^Division of Infectious Diseases & Geographic Medicine, Department of Medicine, Stanford University School of Medicine; Stanford, CA, USA.

^44^Department of Chemical and Systems Biology, Stanford University School of Medicine, Stanford, CA, USA

^45^Division of Nephrology, Stanford University School of Medicine; Stanford, CA, USA.

^46^Veterans Affairs Palo Alto Health Care System; Palo Alto, CA, USA.

^47^Vera Moulton Wall Center for Pulmonary and Vascular Disease, Stanford University School of Medicine; Stanford, CA, USA.

^48^Department of Urology, Stanford University School of Medicine, Stanford, CA, USA

^49^Division of Hematology/Oncology, Department of Medicine, San Francisco Veterans Affairs Health Care System, San Francisco, CA, USA.

^50^Department of Biochemistry, University of California San Francisco; San Francisco, CA, USA.

## Supplementary Material

Materials and Methods

Figs. S1 to S16

Tables S1 to S16

References (40–95)

**Resources**

Data portal for Tabula Sapiens (*14*)

Code for the analysis (*36*)

Single cell gene counts and metadata (*37*)

Histology images (*38*)

SpliZ scores (*39*)

**Fig. S1.**
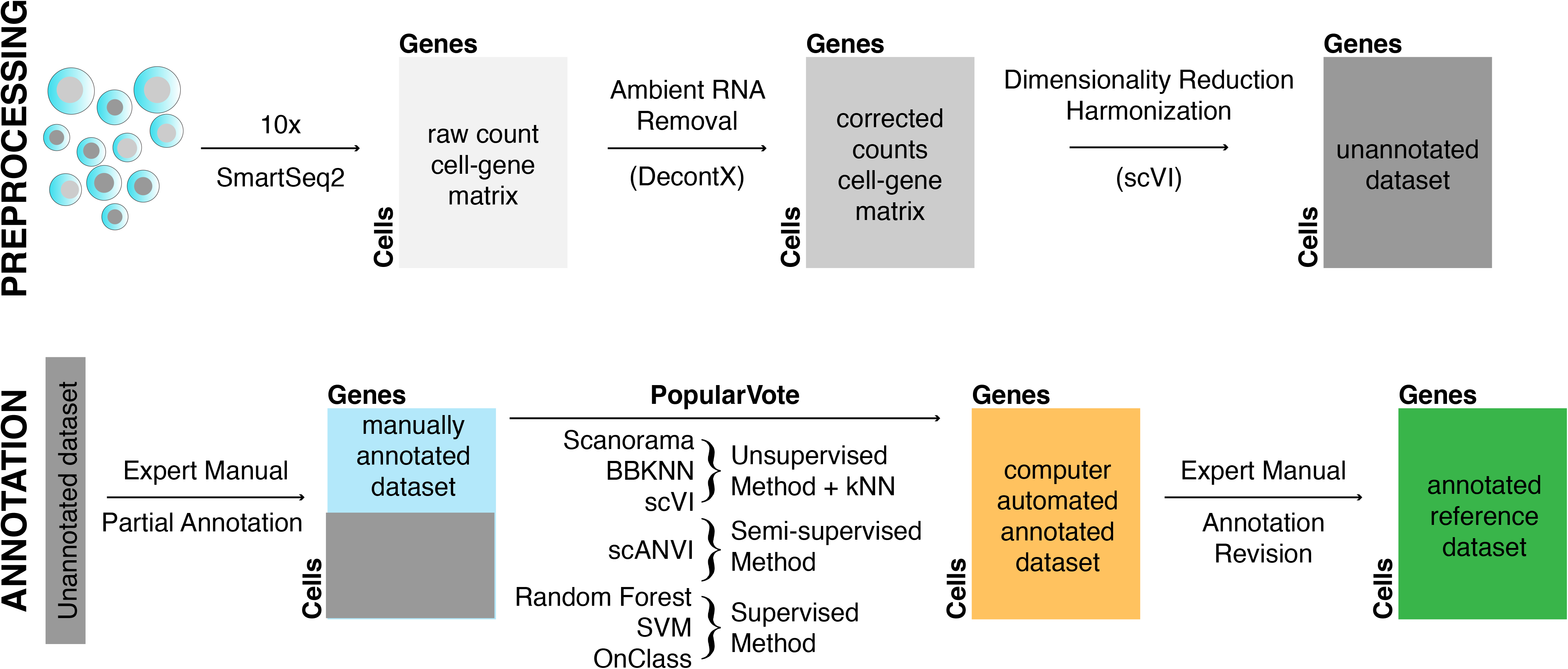

**Fig. S2.**
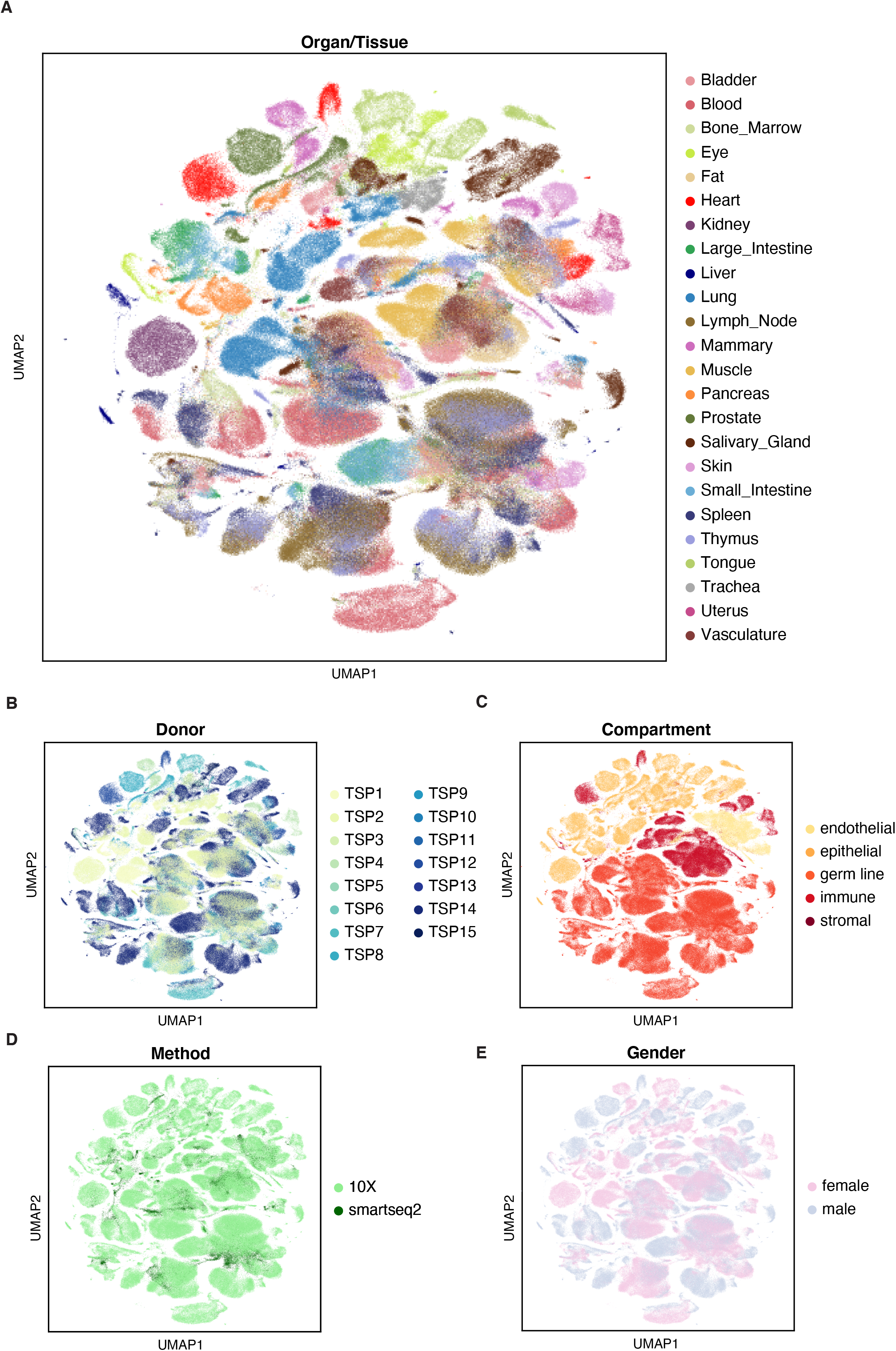

**Fig. S3.**
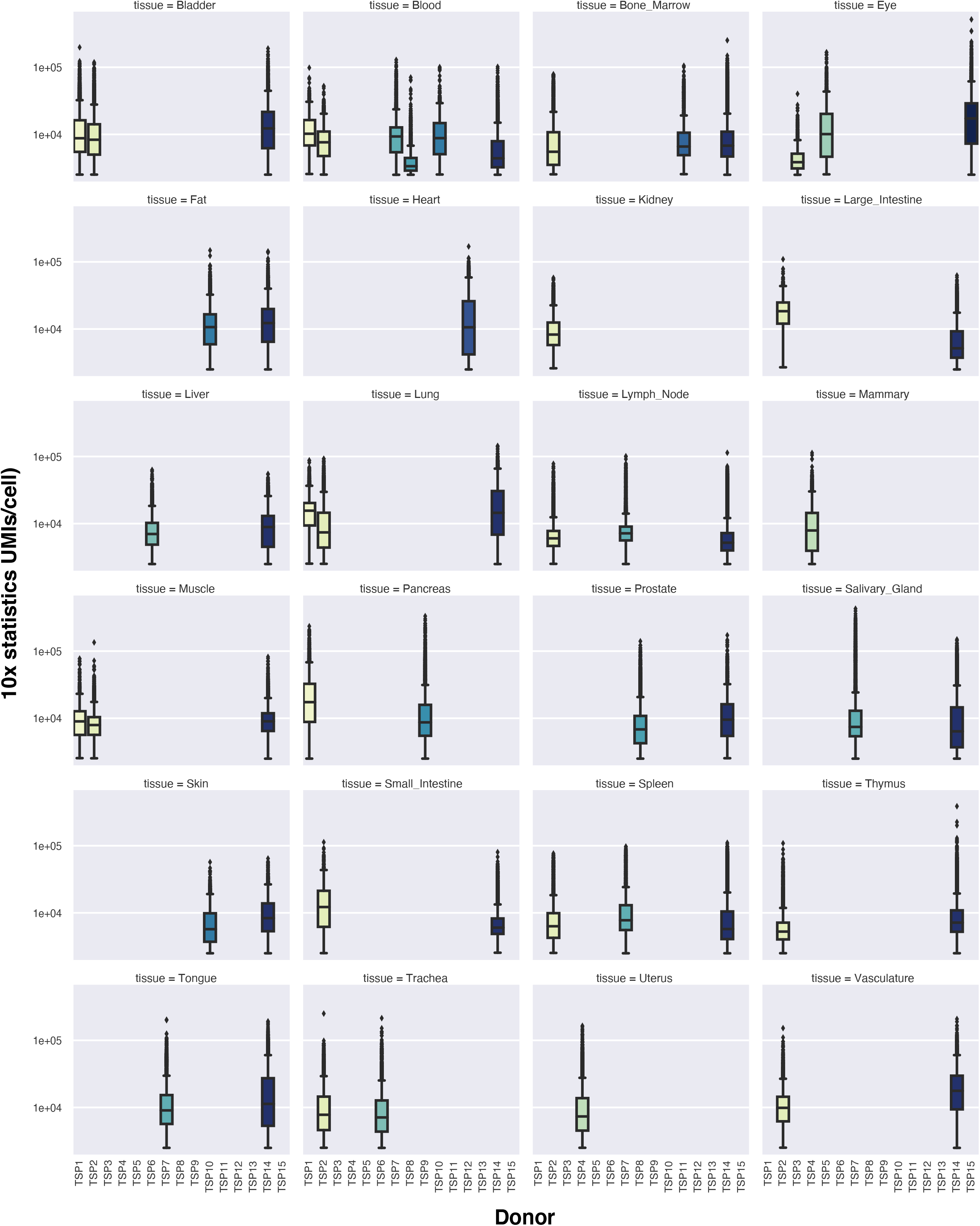

**Fig. S4.**
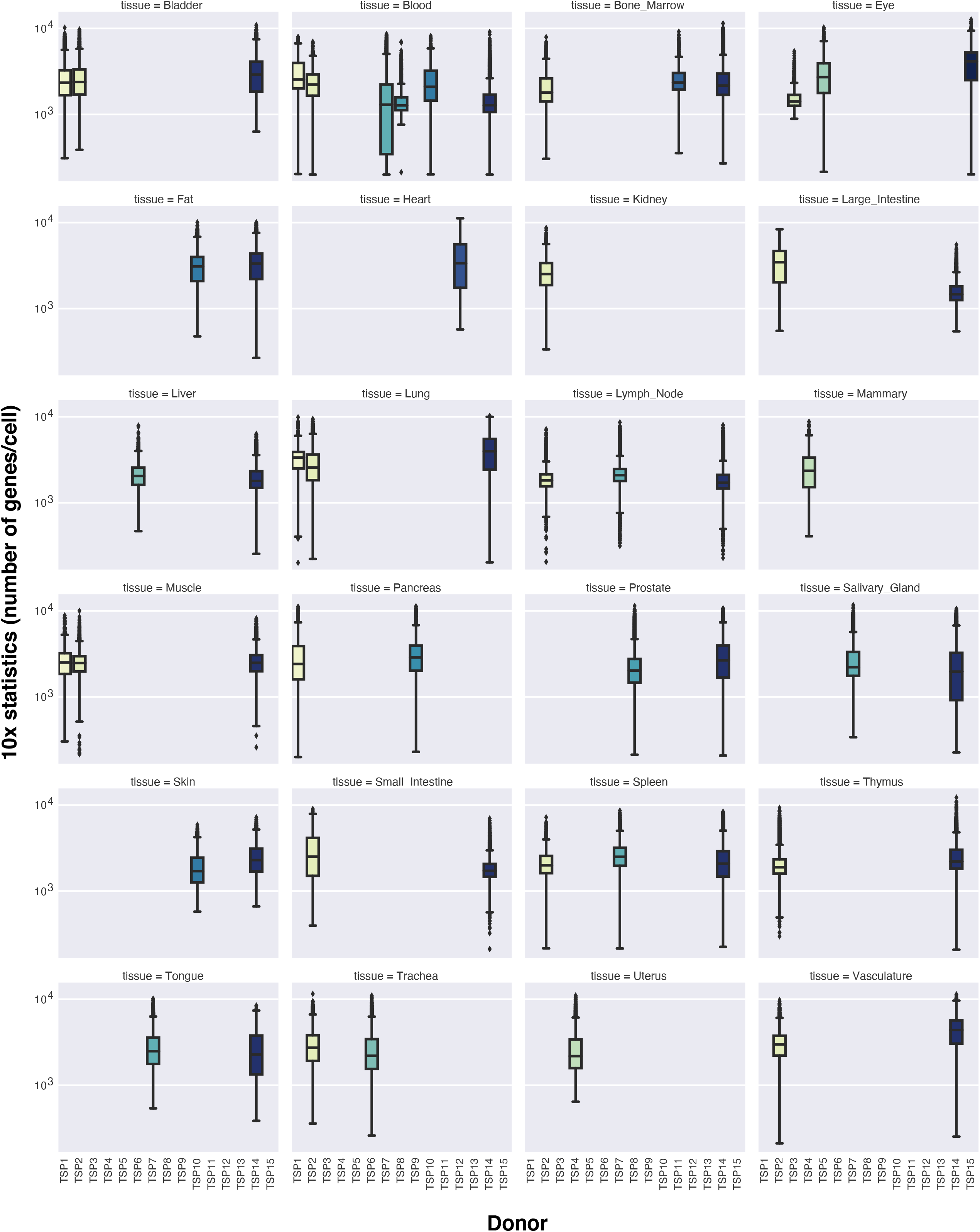

**Fig. S5.**
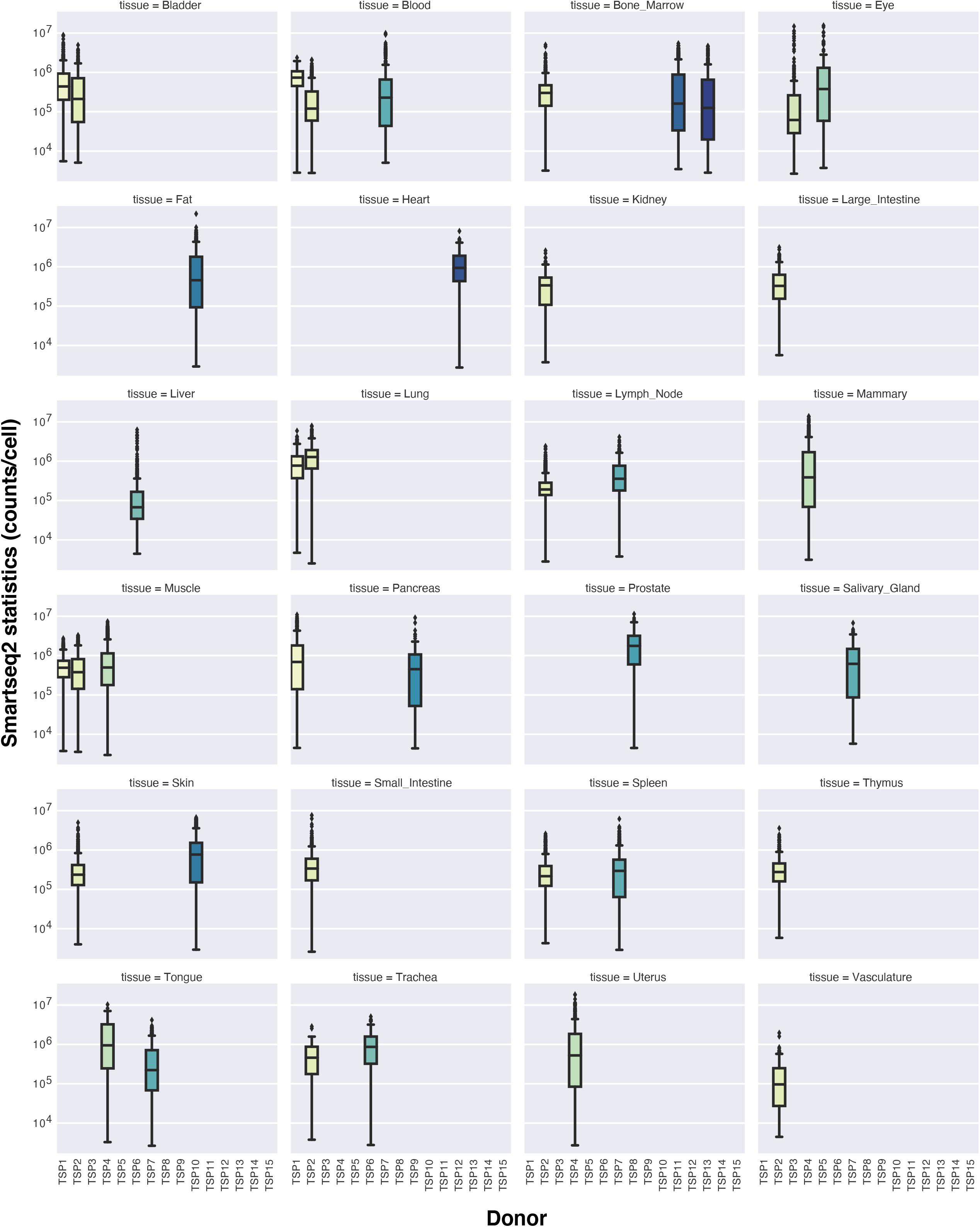

**Fig. S6.**
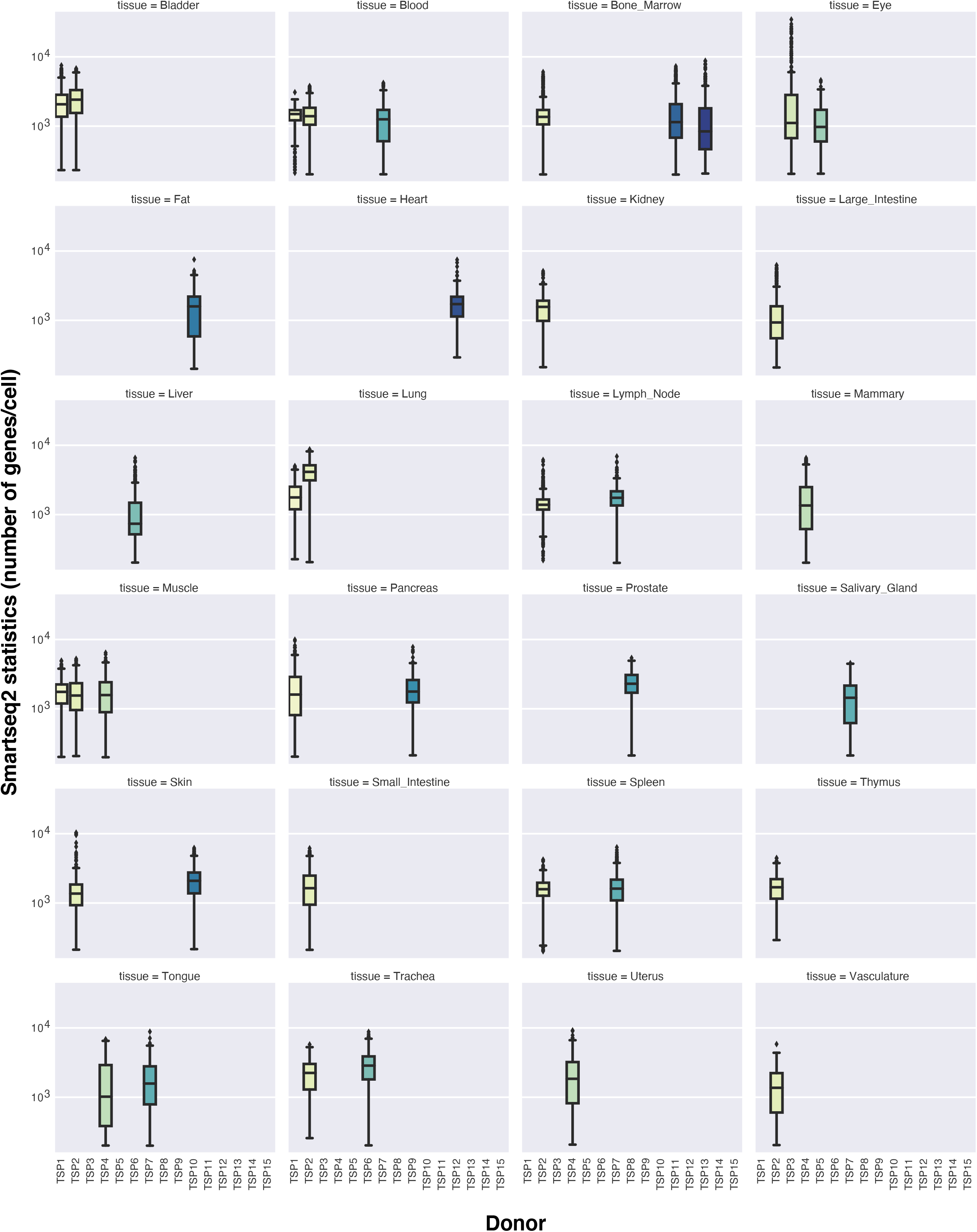

**Fig. S7.**
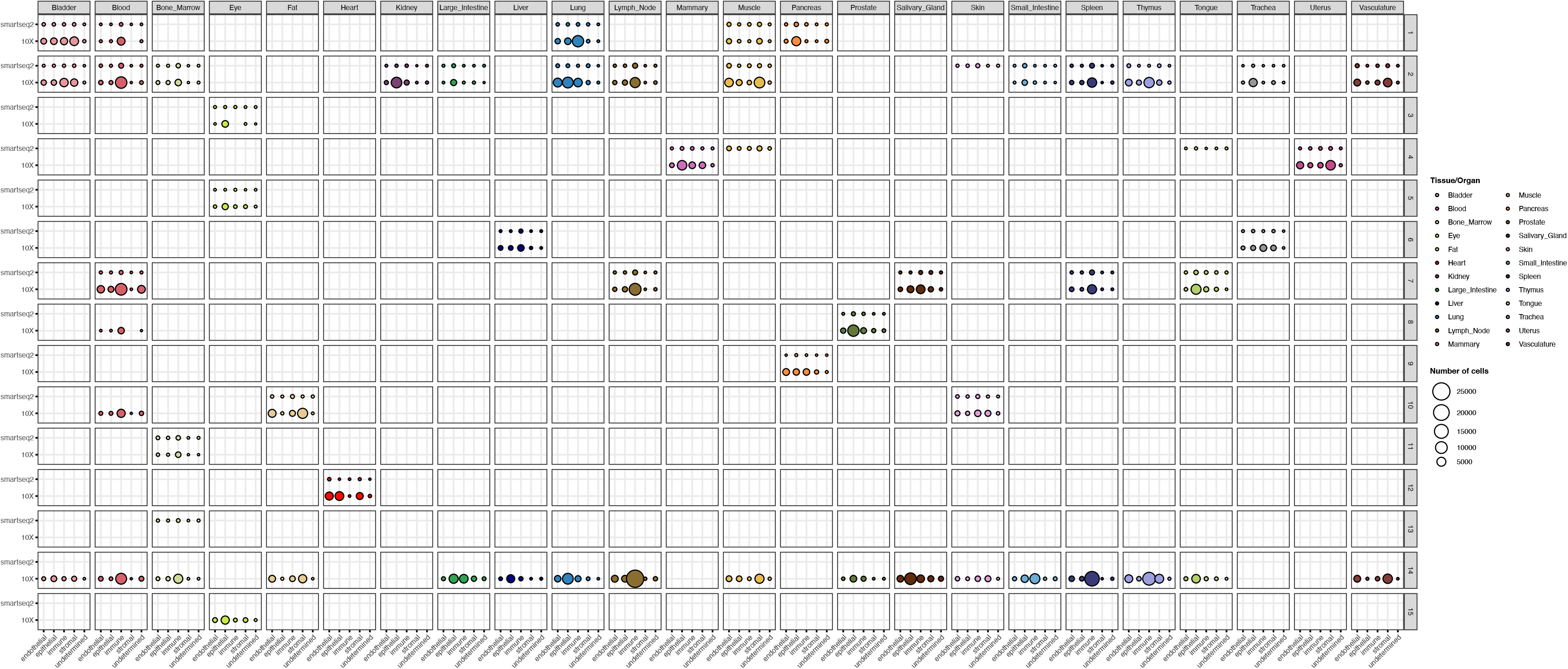

**Fig. S8.**
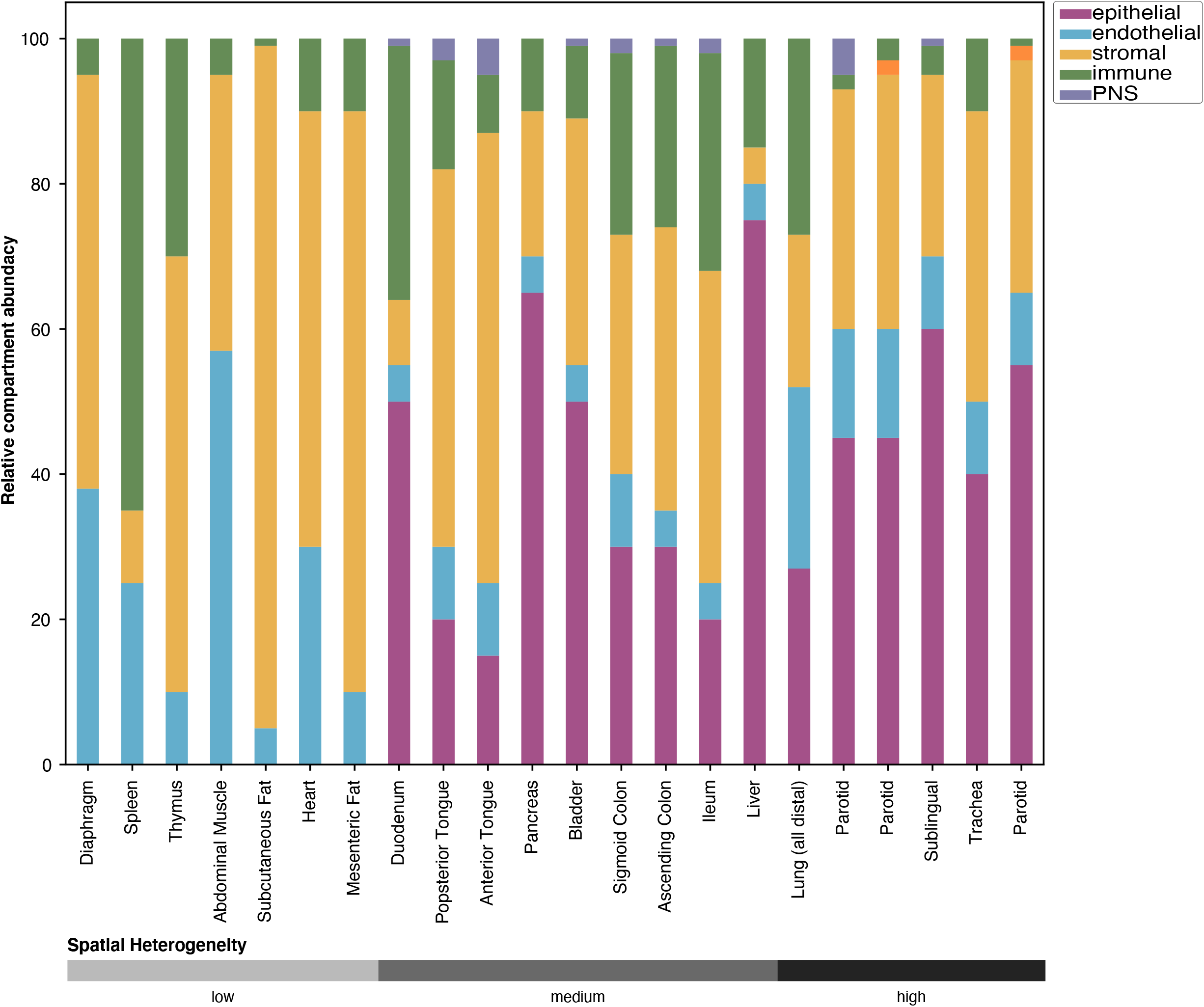

**Fig. S9.**
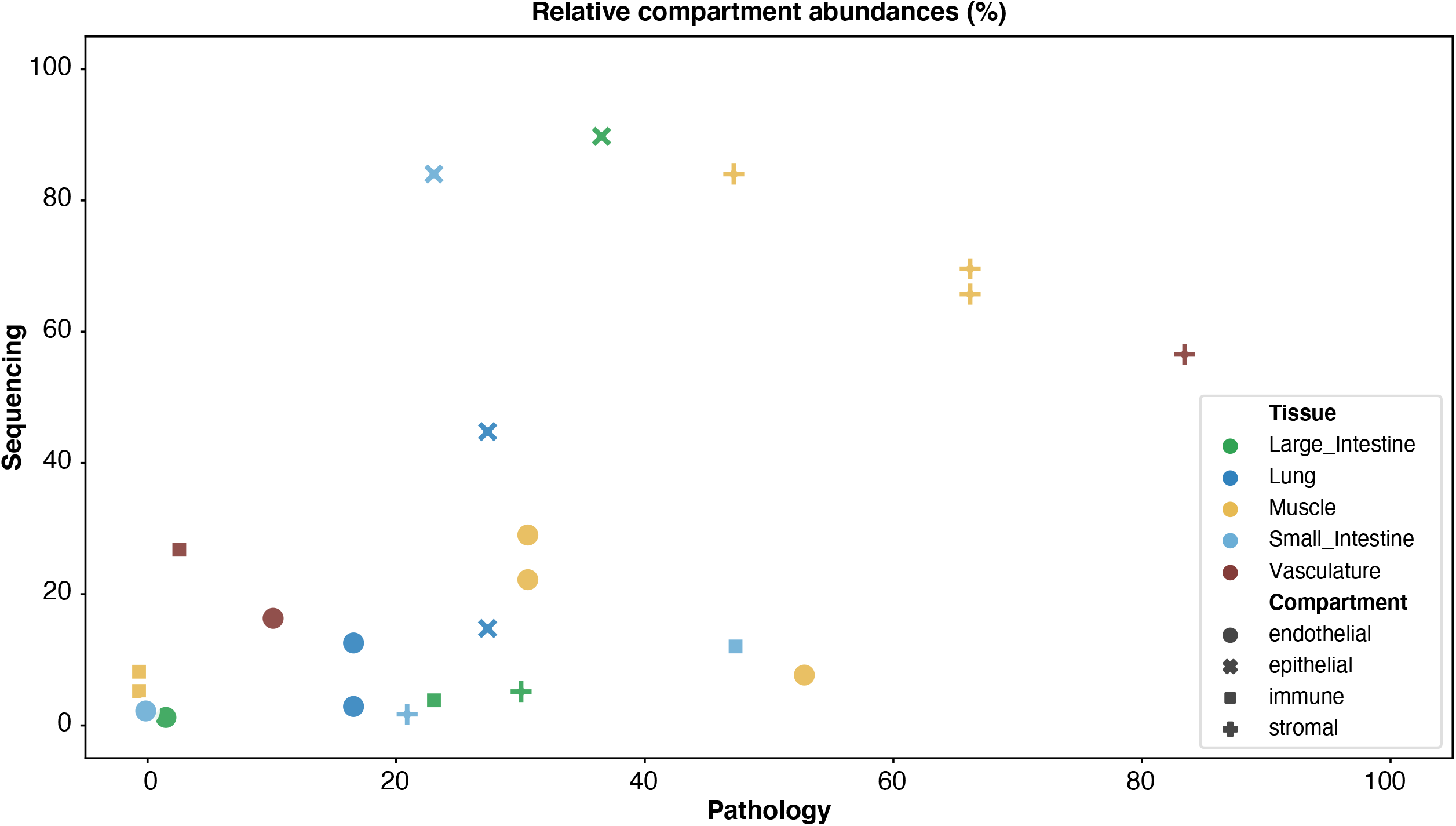

**Fig. S10.**
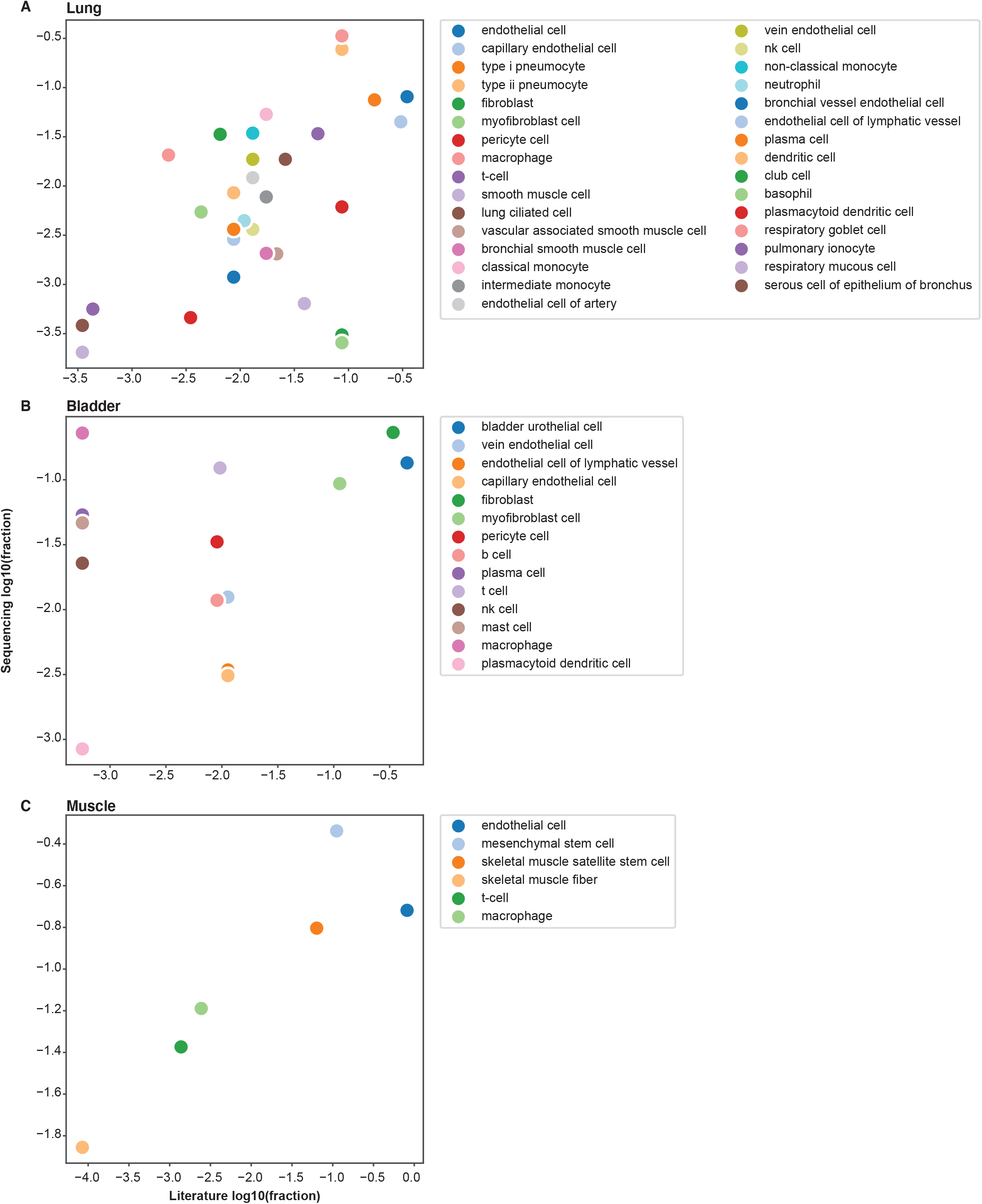

**Fig. S11.**
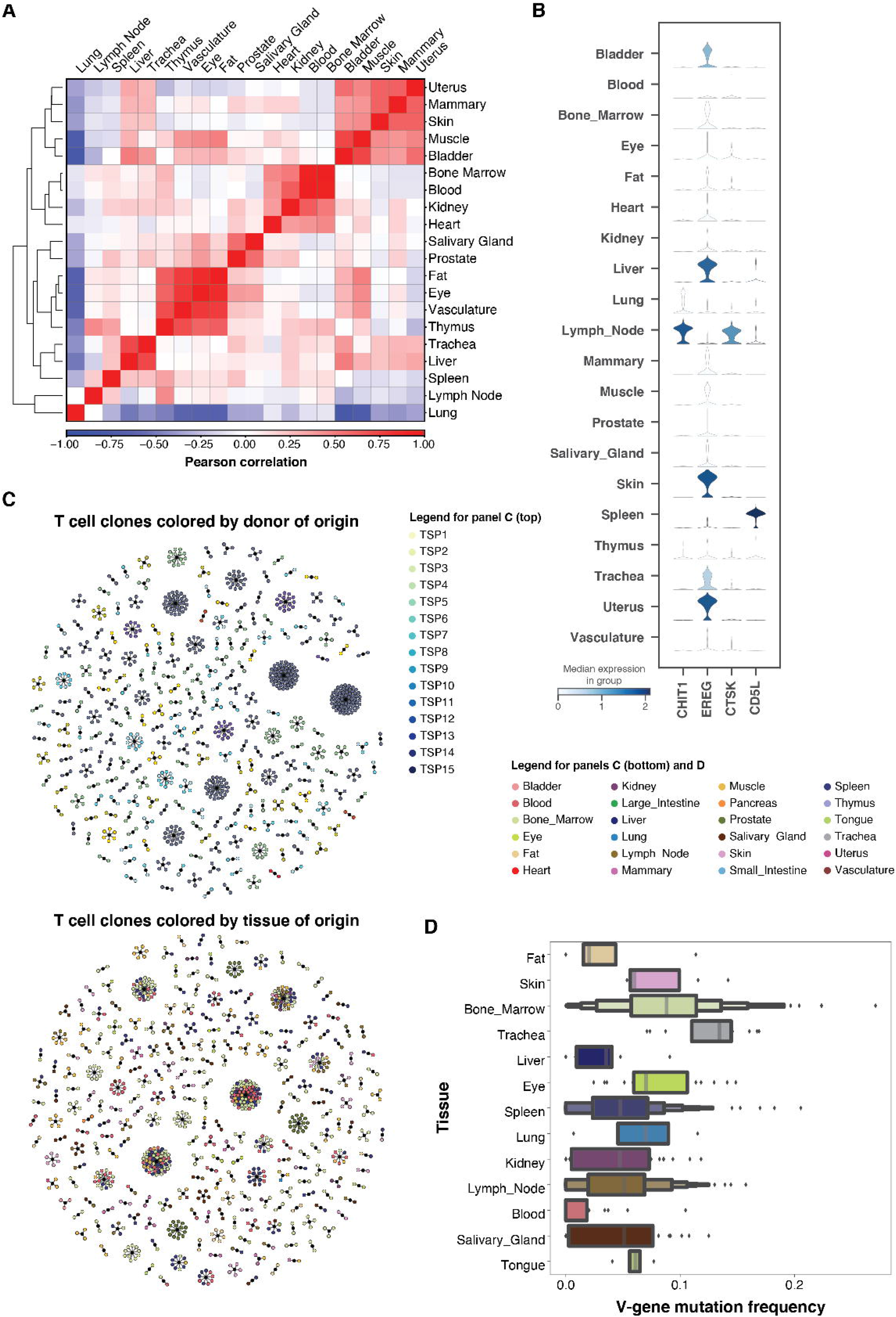

**Fig. S12.**
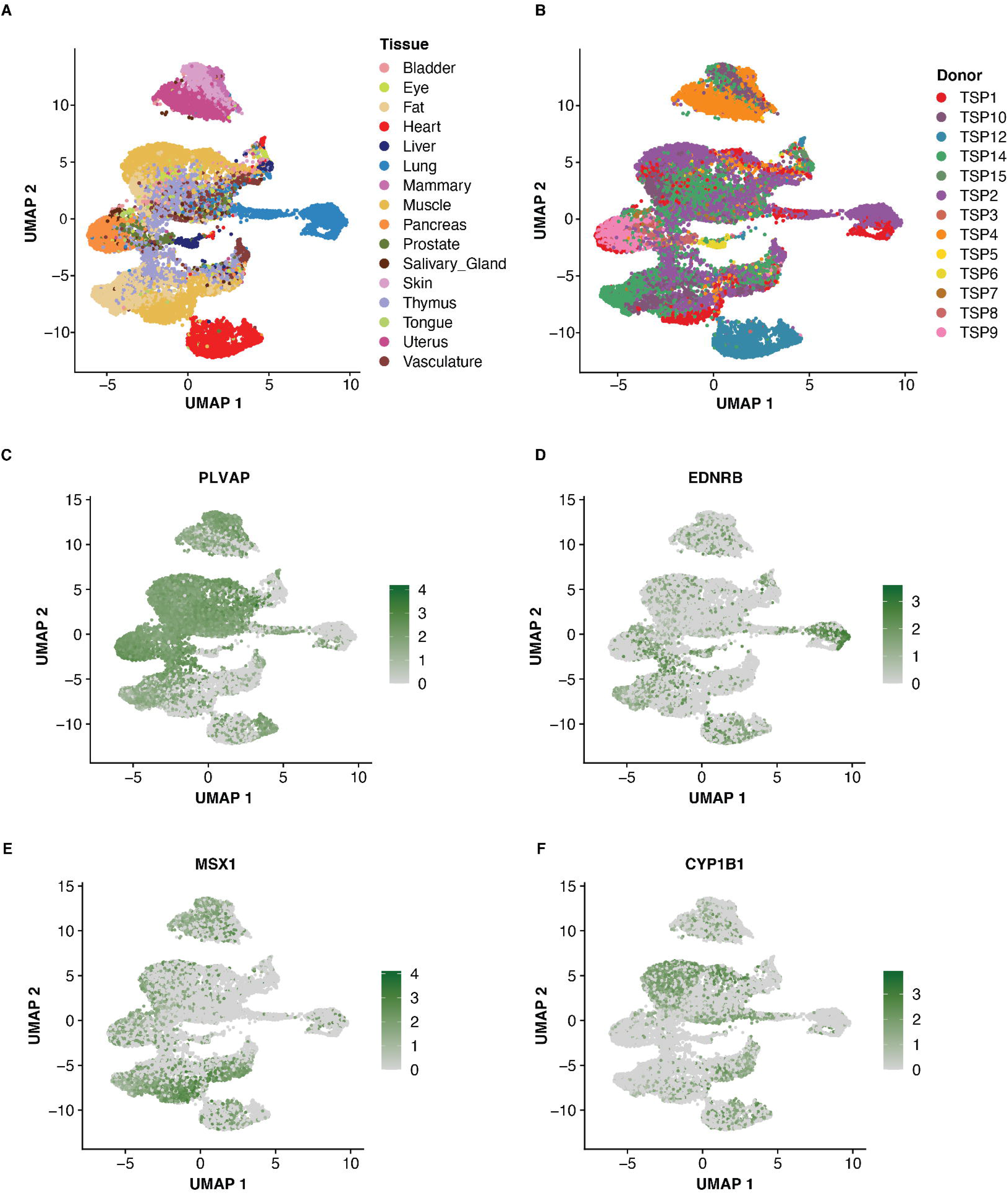

**Fig. S13.**
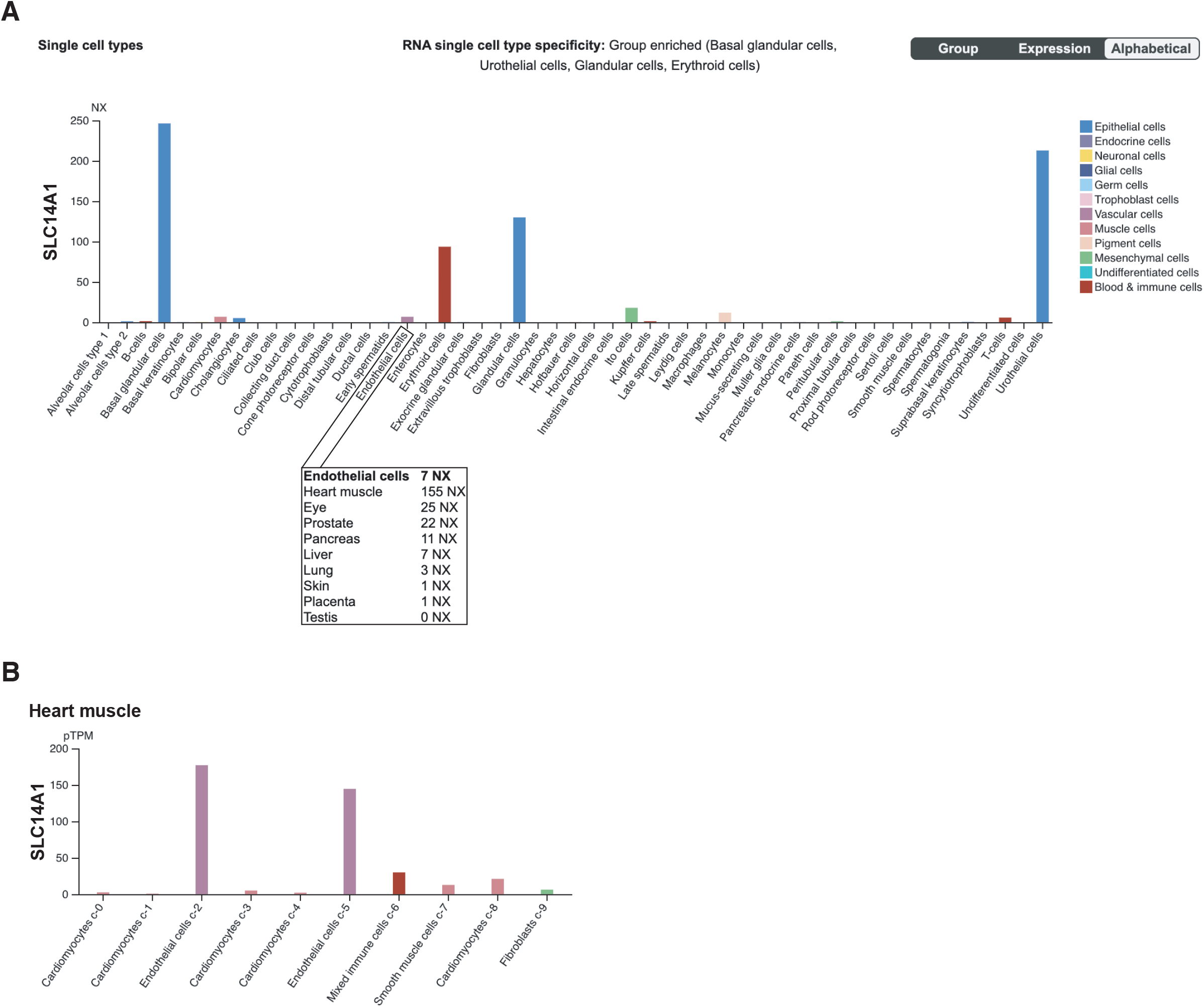

**Fig. S14.**
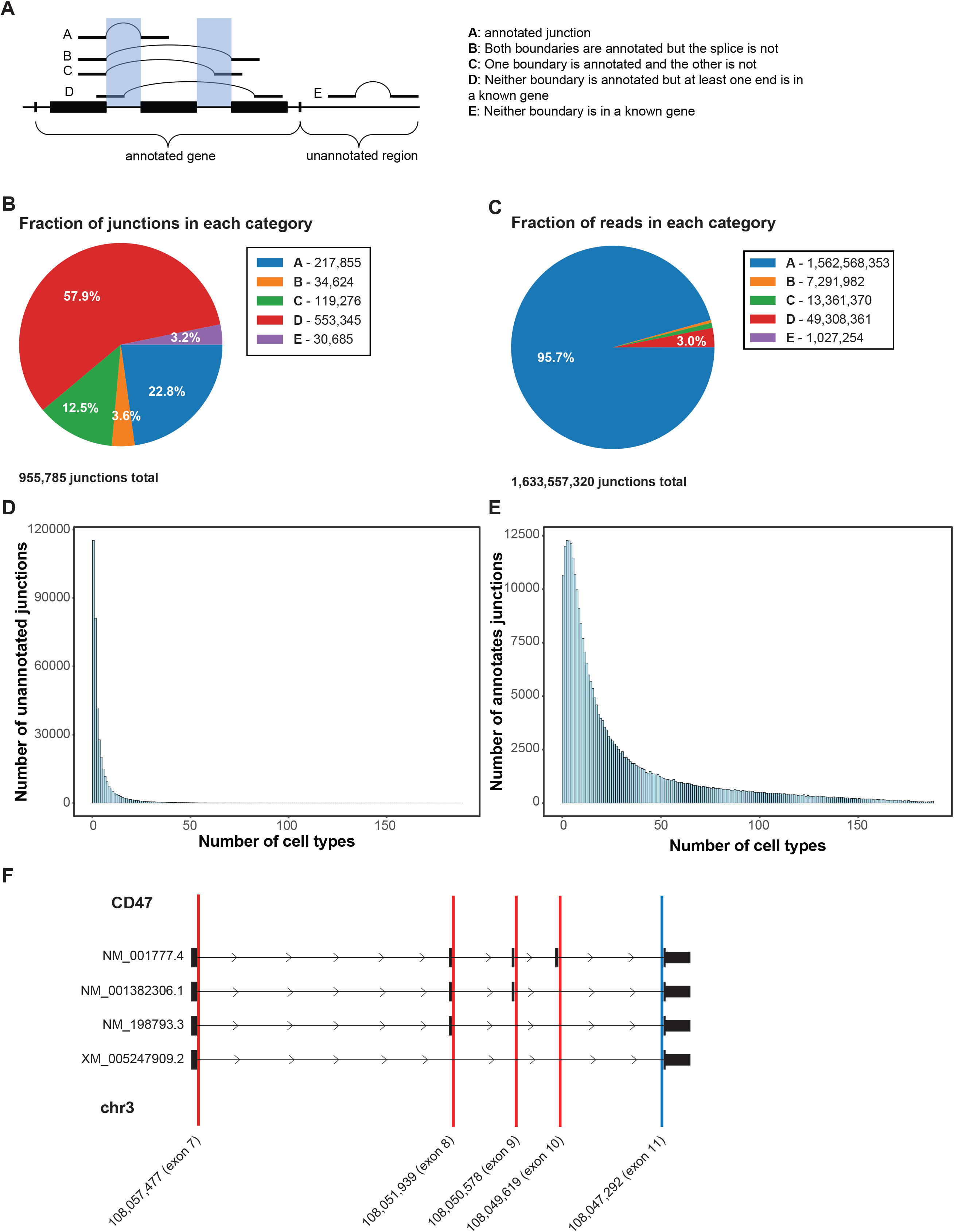

**Fig. S15.**
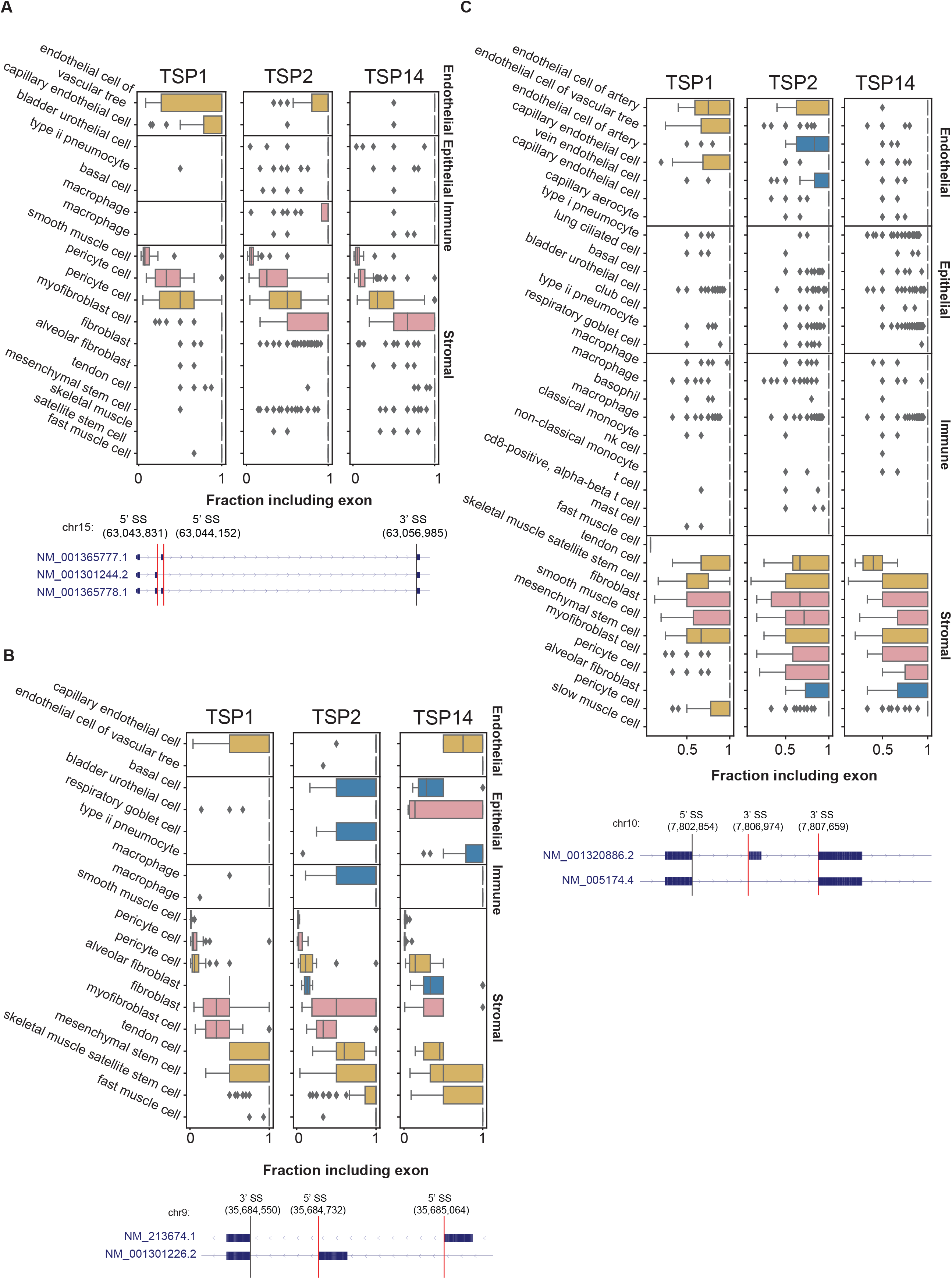

**Fig. S16.**
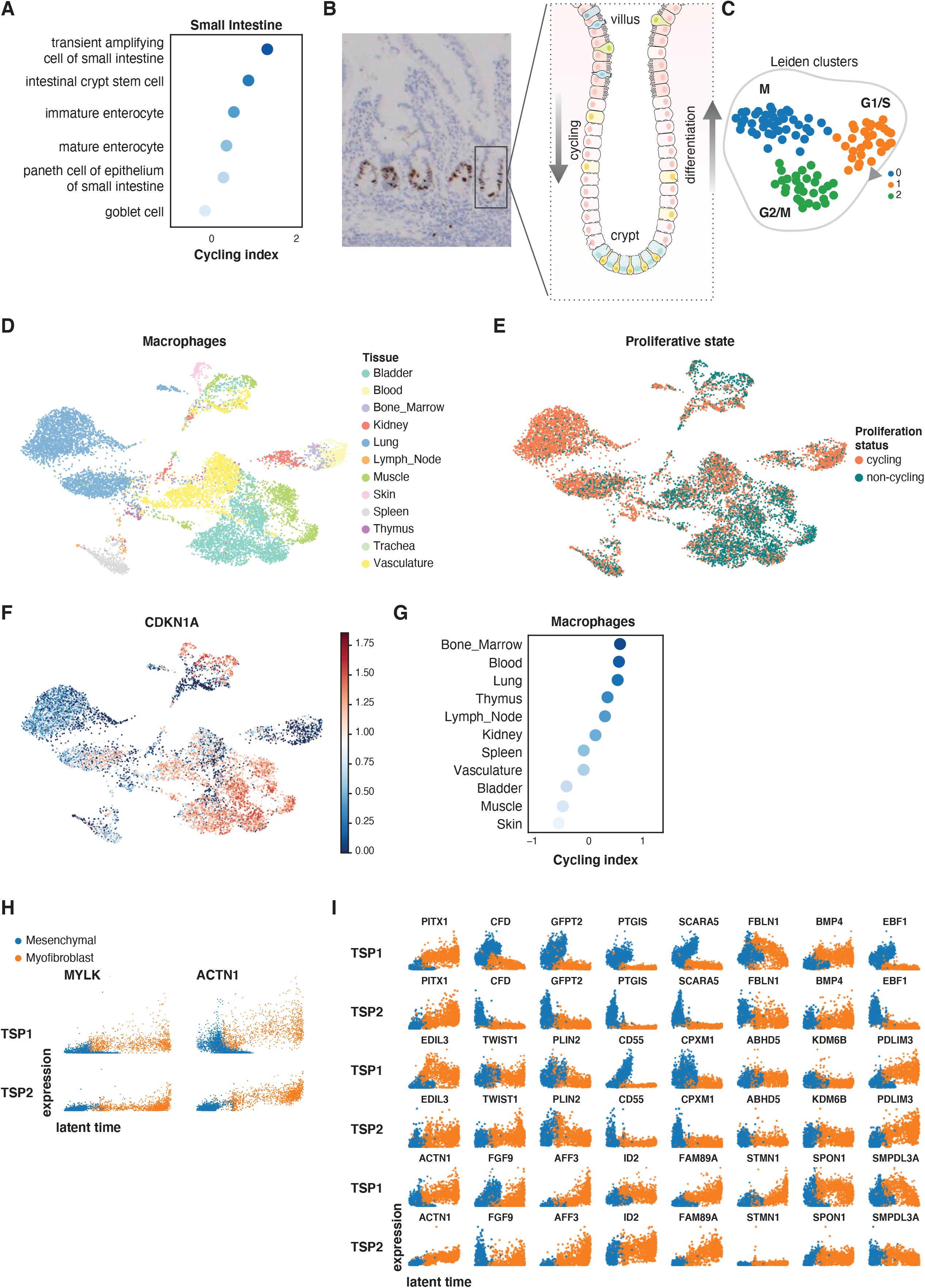

**Fig. S17.**
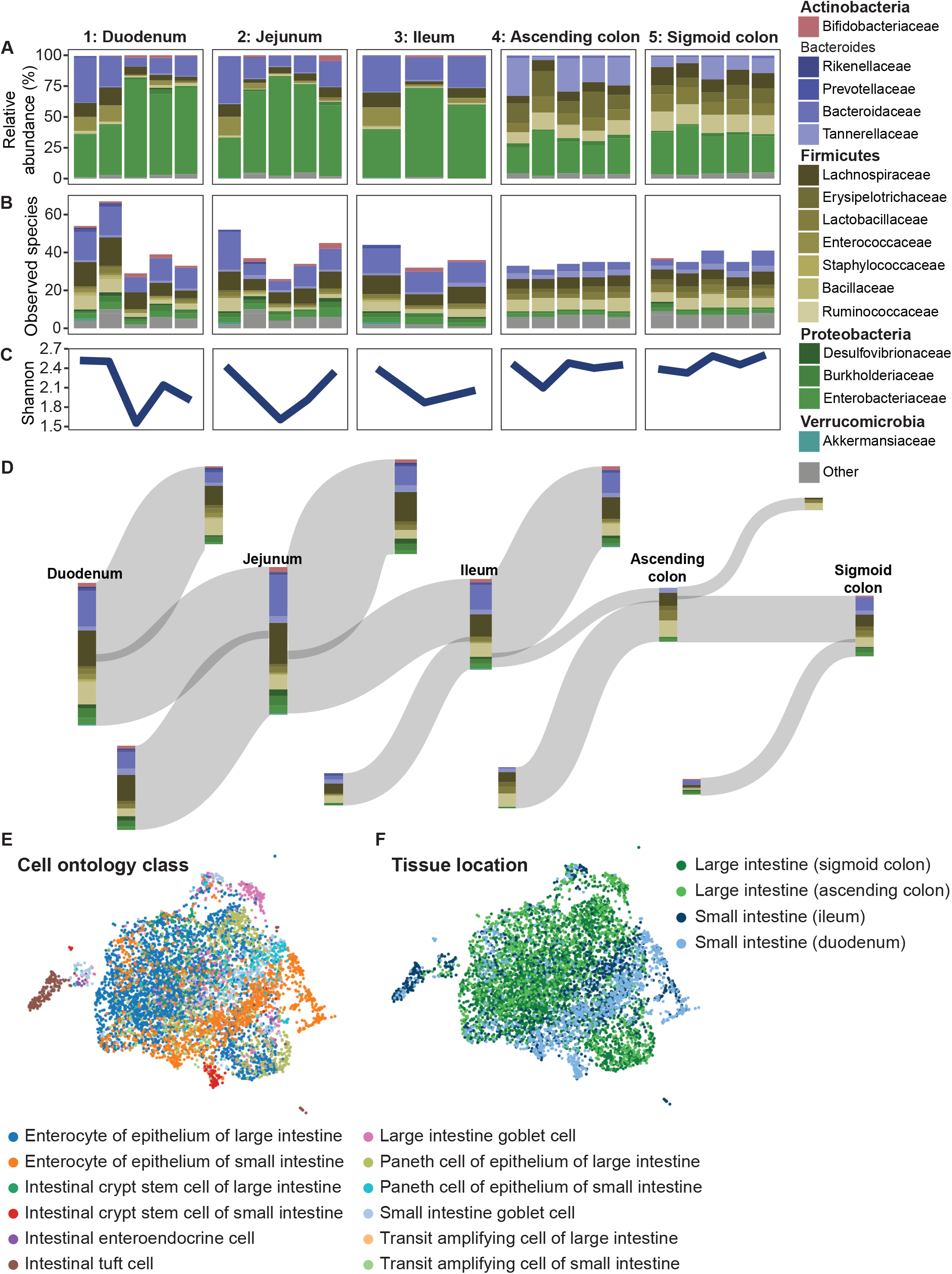

